# Physiological and Molecular Responses of Projected Future Temperatures on Potato Tuberization

**DOI:** 10.1101/2024.08.02.606361

**Authors:** Abigail M. Guillemette, Guillian Hernández Casanova, John P. Hamilton, Eva Pokorná, Petre I. Dobrev, Václav Motyka, Aaron M. Rashotte, Courtney P. Leisner

## Abstract

Potato (*Solanum tuberosum* L.) is one of the most important food crops globally and is especially vulnerable to heat stress. Significant knowledge gaps remain however, in our understanding of the developmental mechanisms associated with tuber responses to heat stress. This study uses whole-plant physiology, transcriptomics, and hormone profiling to gain insights into the mechanisms associated with heat stress impacts on potato tuber development. When plants were grown in projected future temperature conditions, levels of abscisic acid (ABA) were significantly decreased in leaf and tuber tissues while rates of leaf carbon assimilation and stomatal conductance were not significantly affected. While plants grown in elevated temperature conditions initiated more tubers on average per plant, there was a significant decrease (66%) in mature tubers at final harvest. We hypothesize that reduced tuber yields at elevated temperatures are not due to reductions in tuber initiation, but due to impaired tuber filling. Transcriptomic analysis found significant changes in transcript expression for genes related to response to ABA, heat and auxin biosynthetic process. The known tuberization repressor genes SELF PRUNING 5G (*StSP5G*) and CONSTANS-LIKE1 (*StCOL1*) were found to be differentially expressed in tubers grown in elevated temperatures. IDENTITY OF TUBER 1 (*StIT1*) and TIMING OF CAB EXPRESSION 1 (*StTOC1*) are other known tuberization genes that displayed distinct expression patterns in elevated versus ambient temperatures but were not differentially expressed. This work highlights potential gene targets and key developmental stages associated with tuberization to development more heat tolerant potatoes.

## Introduction

Since the industrial revolution, average global temperatures have risen by approximately 1.1°C, and unless carbon dioxide (CO_2_) emissions are substantially diminished, global surface temperatures are expected to continue to increase by an additional 1.5-8°C by next century (IPCC, 2021). These higher-than-optimum temperatures have already been shown to negatively affect productivity and yield in many major crop plants, including potato (*Solanum tuberosum* L.) (Dahal et al., 2019). Potatoes are the fourth most important food crop globally with a total production of 374 million metric tons in 2022 (FAO, 2022). It is estimated that climate change, including temperature fluctuations and heat stress, has the potential to reduce potato production by up to 18-32% by mid-century (Hijmans, 2003). As a staple crop in many countries, understanding how elevated temperatures associated with climate change affect potato development and yield are of utmost importance for future global food security (de Haan & Rodriguez, 2016).

Diploid progenitors of modern tetraploid potatoes are native to the more temperate climate of the Andes Mountains of South American (Ortiz, 2001), with optimal temperatures for tuber growth being 14-22°C (Van Dam et al., 1996). Potato plants grown under temperatures even moderately higher than this range (≥ 25°C) face a shift in biomass allocation towards the aboveground plant, leading to longer stems, more leaves, and decreased tuber yield (Hancock et al., 2014). Although the degree of this response varies among potato cultivars, even relatively heat-tolerant cultivars can experience significant yield loss. Additionally, yield loss is exacerbated the earlier in development the heat stress occurs (Rykaczewska, 2013; 2015). There is a significant knowledge gap however, surrounding the mechanism by which elevated temperatures effect tuberization signaling, tuber development, and in turn, potato yield.

Potato tubers develop from a stolon, or underground stem, originating from axillary buds at the base of the main stem. The development of a stolon into a tuber is caused by changes in the carbohydrate metabolism of the plant in response to environmental cues such as photoperiod and temperature (Kondhare et al., 2021). These cues cause a cascade of internal signals controlling tuberization, and the process is characterized by three separate stages: tuber initiation, tuber filling, and maturation (Obidiegwu et al., 2015). During tuber initiation, the stolon apical tip ceases elongation and begins radial cell expansion and division, driven by transport and deposition of sucrose synthesized in the leaves during photosynthesis (Zierer et al., 2021).

During the subsequent filling and maturation stage, the deposition of sucrose and other metabolites in the stolon continues until the tuber reaches maturity.

Tuberization is mediated by complex interactions between molecular tuberization signals and environmental cues (Dutt et al., 2017). The most well studied gene responsible for regulating tuberization is *S. tuberosum* SELF-PRUNING 6A (*StSP6A*), a FLOWERING LOCUS T (FT) homolog and mobile protein that induces sucrose transport from the leaves to tubers (Navarro et al., 2011). During tuber initiation, StSP6A forms the tuberigen activation complex (TAC) in the stolon tips with FD-like proteins StFDL1a and StFDL1b, which are basic leucine zipper (bZIP) transcription factors (Teo et al., 2017). *StSP6A* is part of the PHOSPHATIDYLETHANOLAMINE BINDING PROTEINS (PEBP) family along with 14 other genes related to flowering/tuberization (Zhang et al., 2022). Under heat stress, it is known that during early stages of tuberization, *StSP6A* is inhibited post-transcriptionally through RNA-based interference by a small RNA called SUPPRESSING EXPRESSION OF SP6A (*SES*), while transcriptional regulation of *StSP6A* is the main regulation process in later stages of tuber development under heat stress (Lehretz et al., 2019; Park et al., 2022). Studies using *StSP6A* over-expression lines found increased tuber count under high temperatures compared to wild type (WT) plants, but yield/tuber weight was still significantly decreased compared to WT in ambient temperature (Park et al., 2022). This indicates that while *StSP6A* plays a significant role in tuber formation, other genes are responsible for the continued growth and development of tubers to maturity when grown in elevated temperatures.

The *StCOL1-StSP5G* inhibitory pathway is known to be an important suppressor of tuberization. Under long-day photoperiods, the StCOL1 protein accumulates and in turn activates *StSP5G*, an inhibitor of *StSP6A* (Abelenda et al. 2016). Both genes are known to be expressed primarily in leaves, although their expression has also been found in tubers (Park et al., 2022).

Another inhibitor of *StSP6A* is TIMING OF CAB EXPRESSION1 (TOC1), a circadian clock gene known to interact with the promoter of *StSP6A*. Studies have shown that *TOC1* has increased expression in elevated temperature and interacts directly with the StSP6A tuberization signal, suppressing its positive feed-forward regulation in stolons (Morris et al.,2019).

Despite recent developments in understanding the effects of heat stress on potato yield, the exact molecular mechanisms controlling tuberization under elevated temperatures remain unknown. Furthermore, few studies have observed developing tubers under chronic heat stress that mimic realistic projected future climate conditions. To address this knowledge gap, we employed dynamically down-scaled global climate projections to determine anticipated future growth conditions for a major potato-production region for the mid-21^st^ century to study the effects of elevated temperature on potato development. In this controlled growth chamber experiment, we investigated the physiological and molecular changes of both source and sink tissue over time under ambient and elevated temperature conditions. Findings from this study have the potential to advance our understanding of the physiological mechanisms associated with potato tuberization in elevated temperature and identify key tuberization genes that may help enhance potato resilience to future elevated temperatures.

## Materials and Methods

### Growth Chamber Experimental Design

Potato plants were grown from seed tubers of the chip-processing cultivar Manistee in a controlled growth chamber experiment. In 2021, seed tubers were obtained from Michigan State University, courtesy of Dr. Dave Douches, and stored at room temperature for 3 weeks to break dormancy. Sprouted seed tubers were cut into halves and planted in 14.2 L pots with Promix BX soil. Plants were grown using 60% relative humidity in controlled growth chambers (Conviron Adaptis; Controlled Environments, Inc) under either ambient (AmbT) or elevated temperature (ElevT) conditions. AmbT conditions represented the average temperature (minimum and maximum) from 1980-2000 (the established control climate period) while ElevT conditions were those projected for the mid-21st century (2040-2060) in a major potato production region in the United States (Eau Claire, Michigan) (Leisner et al. 2017) (Supplementary Table 1).

Future projections of surface minimum and maximum temperature for the ElevT growth conditions were obtained from an ensemble of a dynamically downscaled climate simulations produced by the North American Regional Climate Change Assessment Program (NARCCAP) (Mearns et al., 2012). The climate projections are developed from a combination of regional climate model (RCM) and global climate model (GCM) simulations. The RCM/GCM combination used for this experiment was the Canadian Regional Climate Model (CRCM)/Canadian Global Climate Model version 3 (CGCM3), thereafter referred to as CRCM_cgcm3. The climate model used in this experiment (CRCM_cgcm3), projects that by the mid-21^st^ century the largest change in maximum and minimum temperature in Michigan will occur during tuber development (initiation and filling) (Figure 1) and has previously been observed to impact tuber yield (for detailed methodology on the global climate model downscaling see Leisner et al., 2017).

**Figure 1.**
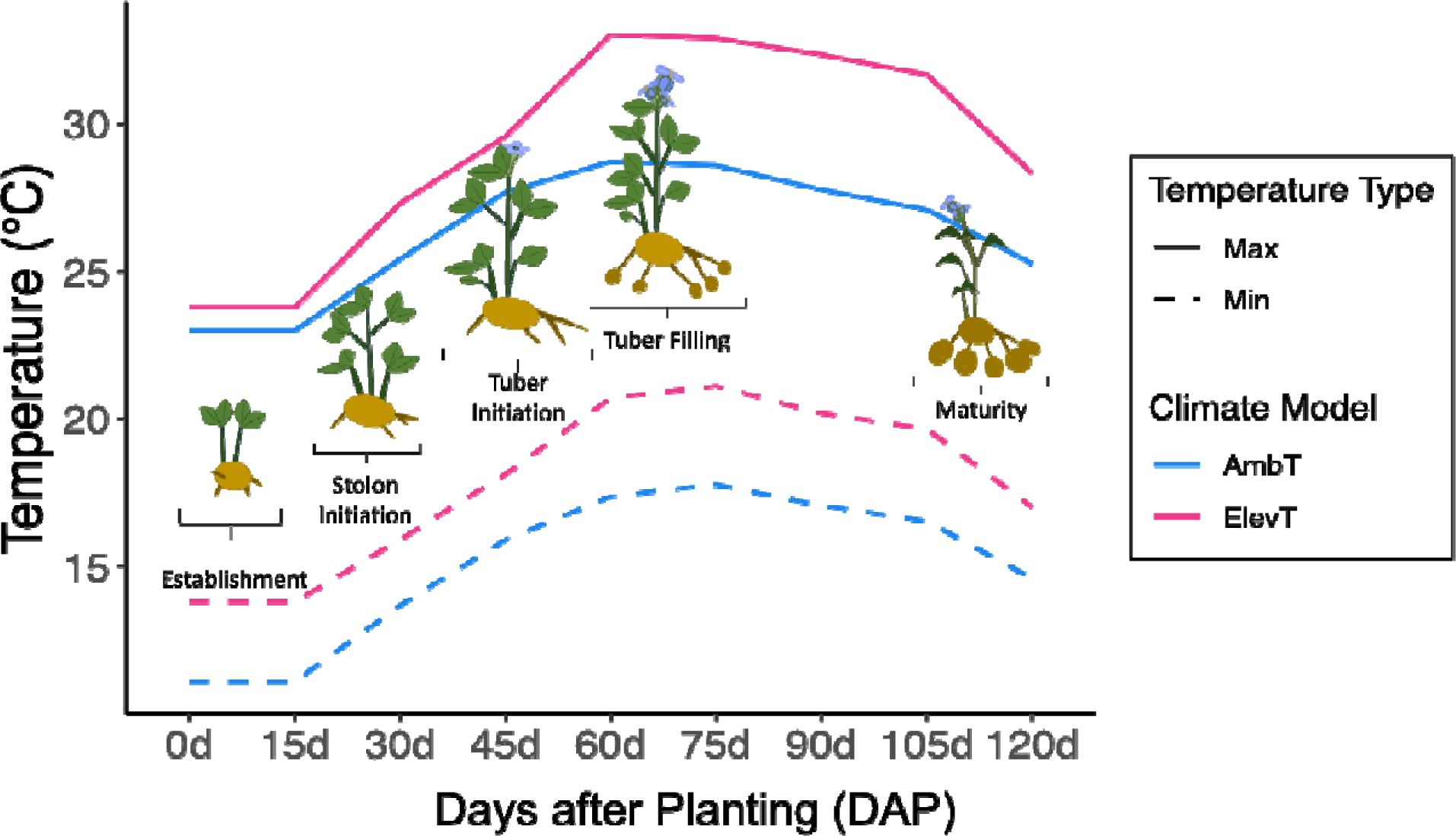
Maximum and minimum temperatures for the climate control period (1980-2000) and projected future temperature for the mid-century (2040-2060) using the CRCM_cgcm3 climate model with plant development stages superimposed on the temperature values. Temperatures are given for the 120-day growing period for potato. CRCM_cgcm3 data obtained from Leisner et al., (2017).

Minimum and maximum temperatures [AmbT: 11.1 - 28.8°C; ElevT: 13.8 - 33°C] along with photoperiod changed every two weeks to mimic seasonal fluctuations during a normal potato growing season (mid-May to mid-September) (Figure 1; Supplementary Table 1). The light intensity in the growth chambers was measured at the top of the canopy using a LI-180 spectrometer (LICOR Biosciences, Lincoln, NE, USA) and ranged from 300-500 μmol m^−2^ s^−1^, and the height of the light rack was adjusted every 2 weeks to maintain these light levels as plants grew. Plants were watered twice a week with 1L of water per plant until 60d, after which they were watered 2-3 times a week to maintain similar soil moisture. Each chamber was fertilized with ∼14g Jack’s Classic All Purpose 20-20-20 Fertilizer diluted in 7L of water once a week once sprouts emerged, about two weeks after planting (JR Peters Inc., Allentown, PA, USA). The plants were rotated twice a week across chambers of the same treatment to minimize chamber effects. Sampling was done on 2 - 4 plants of each treatment every 30 days (d) after planting until maturity (120d). After sampling the plants were removed from the chamber.

Samples taken from plants within the same chamber at the same time point were pooled together as technical replicates, and each chamber was used as a biological replicate (*n* = 2).

### Photosynthetic Gas Exchange, Chlorophyll Fluorescence, and Chlorophyll Content Measurements

Midday gas exchange measurements of one leaf per plant was recorded *in situ* between 11:00 and 13:00 to determine photosynthetic CO_2_ assimilation (*A*) and stomatal conductance (*g_s_*) using a LI-6800 portable photosynthesis system (LICOR Biosciences, Lincoln, NE, USA). Leaf temperatures set to reflect chamber settings (16.5 and 18.7°C at 30d, 20.1 and 23.8°C at 60d, 19.9 and 23.4°C at 90d, and 17.7 and 20.3°C at 120d for AmbT and ElevT temperatures, respectively). Relative humidity, CO_2_ concentration, and light intensity levels were also set to mimic conditions within the growth chambers. Following gas exchange measurements, two fresh leaves from each sampled plant were collected and four, 6mm diameter leaf discs were taken, placed adaxial side up on 3mM MES buffer. Maximum Photosystem II (PSII) quantum efficiency of the leaf discs was measured in terms of F_v_/F_m_, which is variable fluorescence (F_v_) divided by the maximum fluorescence (F_m_), at 24, 48, 72, and 96h after dark exposure as described by Zwack et al. (2016). Changes in F_v_/F_m_ were taken to measure the rate of leaf senescence. After 4 days, each leaf disc was then transferred to 500µL of 100% methanol overnight, and a UV/Vis spectrophotometer was used to measure total chlorophyll of the methanol extract for each leaf disc as described by Zwack et al. (2016) (Beckman Coulter, USA).

### Biomass and Yield Measurements

Aboveground biomass and yield measurements were taken at 120d for four plants per chamber. Plant height was taken by measuring the tallest point of each plant. The entire aboveground plant was cut off, and fresh weight (FW) was recorded before placing the plants in a 60°C dryer for 2 weeks to measure dry weight (DW). Tubers were collected fresh, counted per chamber, and categorized into 3 different developmental classes based on weight: Tuber Initials (TI) (< 0.6g), Intermediate Tubers (IMT) (0.6-5g), and Mature Tubers (MAT) (> 5g). Examples of each tuber size class are given in Supplementary Figure 1. For tuber count and weight measurements at each sampling point, 2 or 4 individual plants were sampled from each condition. Two plants were sampled at 30d and 90d, while four plants were sampled at 60d and 120d. Final tuber yield per chamber (at 120d) was taken by weighing all tubers with a mass greater than 0.6g (TIs were excluded from final yield). Values were averaged across chambers (*n* = 2).

### Phytohormone Extraction and Analysis

Phytohormone analysis was done on leaf samples at 90d and 120d. Tissue samples were collected from one leaf per plant (four plants total) and collected and immediately flash frozen in liquid nitrogen. Tubers were collected and separated into size classes before flash freezing.

Pooled tissue was grounded into a fine powder and stored at -80°C. Between 10-20mg of frozen ground tissue powder were placed into microtubes and lyophilized overnight in a FreeZone 1 freeze dry system at -50°C (LabConco, Kansas City, MO, USA). Phytohormone determination was done in the Laboratory of Hormonal Regulations in Plants, Institute of Experimental Botany of the Czech Academy of Sciences as previously described in Prerostova et al. (2021).

Homogenized samples (ca. 1.5-2mg DW) were extracted with 100µL 1M formic acid solution. Mixtures of stable isotope-labeled phytohormone standards were added at 1pmol per sample. The extracts were centrifuged at 30,000 × g for 25 min at 4°C. The supernatants were applied to SPE Oasis HLB 96-well column plates (10mg/well; Waters, Milford, MA, USA) conditioned with 100µL acetonitrile and 100µL 1M formic acid using Pressure+ 96 manifold (Biotage, Uppsala, Sweden). After washing wells three times with 100µL water, the samples were eluted with 100µL 50% acetonitrile in water. An aliquot of the extract was analyzed on a liquid chromatography/mass spectrometry (LC/MS) system consisting of UHPLC 1290 Infinity II (Agilent, Santa Clara, CA, USA) coupled to 6495 Triple Quadrupole Mass Spectrometer (Agilent, Santa Clara, CA, USA), operating in MRM mode, with quantification by the isotope dilution method.

Internal standards used for the phytohormone analysis were: ^13^C_6_-IAA (Cambridge Isotope Laboratories, Tewksbury, MA, USA); ^2^H_4_-SA (Sigma-Aldrich, St. Louis, MO, USA); ^2^H_3_-PA, ^2^H_3_-DPA (both from NRC-PBI, Saskatoon, Canada); ^2^H_5_-tZ, ^2^H5-tZR, ^2^H5-tZ7G, ^2^H5- tZ9G, ^2^H5-tZOG, ^2^H_5_-tZROG, ^2^H_5_-tZRMP, ^15^N_4_-cZ, ^2^H_3_-DZ, ^2^H_3_-DZR, ^2^H_3_-DZ9G, ^2^H_3_- DZRMP, ^2^H_6_-iP, ^2^H_6_-iPR, ^2^H_6_-iP7G, ^2^H_6_-iP9G, ^2^H_6_-iPRMP, (^2^H_5_)(^15^N_1_)-IAA-Asp, (^2^H_5_)(^15^N_1_)- IAA-Glu, (^2^H_5_)(^15^N_1_)-IAM, ^2^H_6_-ABA, ^2^H_5_-JA, ^2^H_2_-GA1, ^2^H_2_-GA4, ^2^H_2_-GA8, ^2^H_2_-GA12, ^2^H_2_- GA19 (all from Olchemim Ltd., Olomouc, Czech Republic).

Data acquisition and processing was performed with Mass Hunter software B.08 (Agilent, Santa Clara, CA, USA). Three technical replicates per sample were averaged together, and technical replicates with a relative standard deviation higher than 30% were excluded.

Biological replicates were then averaged together and analyzed using an ANOVA.

### RNA Extraction and Sequencing

Total RNA was extracted from the same frozen ground tissue used for the phytohormone analysis. RNA from leaf and tuber tissue collected at each time point was done using the Spectrum Plant Total RNA Extraction kit (Sigma-Aldrich, St. Louis, MO, USA). The RNA was treated with Turbo DNA-free kit, and concentration and integrity were quantified using a NanoDrop Microvolume UV-Vis Spectrophotometer (Thermo Scientific, Wilmington, DE, USA) and 2100 Bioanalyzer (Agilent, Santa Clara, CA, USA). Library preparation and sequencing was performed by Novogene (Novogene Corporation, Inc., Sacramento, CA, USA).

Libraries were sequencing using the Illumina Novaseq 6000 platform producing 150-bp paired-end (PE) reads. A total of 49 libraries were sequenced, with an average of 44,601,280 total reads per library (Supplementary Table 2). Raw sequence data are available for download at the National Center for Biotechnology Information (NCBI) Sequence Read Archive (SRA) under the BioProject ID PRJNA962840.

### Differential Gene Expression Analysis

RNA reads from sequencing were quality trimmed (p > 20) and adapters were removed using *fastp* (v0.23; Chen S., 2023), and quality checks were completed using *fastQC* (v0.11.9; Chen et al., 2018). The reads were aligned to the tetraploid Atlantic potato version 3 (ATL_v3) genome from SpudDB using *HISAT2* (Hoopes et al., 2022; downloaded August 2023 from http://spuddb.uga.edu; *v2.2.1* Kim et al., 2019). Gene count matrices were generated with *featurecounts* (*v2.0.3*; Liao et al., 2014) using the representative high confidence gene models of the ATL_v3 genome. The raw count matrices can be found in Supplemental Data File 1 and served as input for the *DEseq2* (*v1.38.3*; Love et al., 2014) R package in which visualization, modeling, and differential syntelog-specific expression analysis of the libraries was completed. Genes with read counts of zero across all libraries were removed from the analysis. *P*-values were adjusted for false discovery rate using a Bonferroni correction method, and genes with *p*- adjusted values < 0.05 were considered as differentially expressed genes (DEGs). Functional annotations were downloaded from SpudDB (Downloaded August 2023 from http://spuddb.uga.edu). To visualize variation in gene expression across different tissues and treatments, a principal component analysis (PCA) was done to reduce the dimensionality of the data to identify underlying patterns. Normalized gene counts generated from *DESeq2* were used as input for the PCA. PCA figures were generated using *ggplot* in R (version 4.3.1).

### Gene Ontology (GO) Association

Gene ontology (GO) association was done on DEGs from this experiment. GO terms were assigned to the ATL_v3 gene models by searching the protein sequences against the Arabidopsis proteome (TAIR10) using DIAMOND (*v2.1.8l*; Buchfink et al., 2015) with a e-value cutoff of 1e-5. The top match was used to transfer the GO annotation from the TAIR 10 Gene Ontology Annotations. The GO terms were then slimmed using the map2slim from the go-perl package (*v0.15;* http://search.cpan.org/~cmungall/go-perl/) to generate the final set of GO slim terms. The file “ATL_v3.working_models.go_slim.obo.gz” generated from this research has been uploaded to the SpudDB database (http://spuddb.uga.edu/ATL_v3_download.shtml). Additional information regarding all GO terms was obtained from TAIR (The Arabidopsis Information Resource (TAIR), https://www.arabidopsis.org/download/overview on March 2024) to generate a full list of associated GO terms with each DEGs for TI (Supplemental File 2).

### Identifying Genes of Interest in Tuber Development in the Atlantic Potato Genome

BLAST (Altschul et al., 1990) was used to identify previously characterized tuberization genes from the double-monoploid (DM) reference genome (*v6.1*; Pham et al., 2020) in the recently published ATL_v3 genome (Hoopes et al., 2022). SpudDB (http://spuddb.uga.edu/) was used to obtain nucleotide sequences of previously characterized genes in DM and then used as query sequences for a nucleotide BLAST (blastn) against the ATL nucleotide database within SpudDB. Additionally, OrthoFinder (v2.5.4; Emms & Kelly, 2019) was used to confirm orthology between ATL genome genes of interest and DM genome genes of interest. Based on our findings, sequences in the ATL genome were labeled in accordance with their existing DM gene nomenclature, augmented with numeric suffixes denoting different gene loci (i.e., syntelogs).

Syntelogs are homologous genes resulting from a whole-genome duplication event or speciation and are expected to share similar functions. Information regarding the reference DM locus and the corresponding ATL syntelog(s) loci can be found within the Supplementary Table 3. TPM values of the genes of interest were obtained from *Salmon* (v.1.9.0; Patro et al. 2017) using alignment mode quantification with the transcript sequences (cDNA) of the representative high confidence gene models set available for the ATL_v3 genome in SpudDB. From our tissue sampling, we then averaged TPM values for each syntelog at each timepoint and tissue (*n* = 2). Exceptions to the average TPM value are TI at 30d and IMT at 90 and 120D where only one biological replicate is available due to lack of sufficient tissue at these time points. Syntelogs graphed in Figure 7 in the manuscript represent syntelog(s) found to be DEGs or those with highest transcript abundance while all syntelogs from the genes described are in Supplementary Figures 5A-G.

### Statistical analysis of physiology data

Assumptions of normality were validated using the Shapiro-Wilk test and homogeneity of variance was checked using the Levene’s test. T-tests, ANOVA, and Tukey post-hoc tests on all physiology data were completed using the *stats* R package (R Core Team, 2022).

### Data Availability

The RNA-Seq raw sequence data of the 49 samples have been uploaded to the National Center for Biotechnology Information (NCBI) Sequence Read Archive (SRA) under the BioProject ID PRJNA962840. Matrices containing the raw counts from our analysis can be found in Supplemental Data File 1. GO association tables from DEGs in TI at each timepoint can be found in Supplemental Data File 2. Syntelogs identified in this manuscript between DM and ATL genotypes with Orthofinder can be found in Supplemental Data File 3.

## Results

### Elevated temperature impacts on leaf physiology

To understand the effects of elevated temperature on potato leaf physiology, gas exchange of the youngest mature leaves was measured to determine rates of carbon assimilation (*A*) and stomatal conductance (*g_s_*) at each time point. Rates of *A* and *g_s_* were not significantly different between treatment groups except at 60d where the elevated temperature (ElevT) plants had significantly lower rates of both *A* and *g_s_* than ambient temperature (AmbT) plants (Figure 2). Although plants from both AmbT and ElevT treatment groups were initially watered equally, at the 60d sampling date, ElevT plants were noticed to have visibly drier soil than the AmbT plants, after which all plants were watered according to soil moisture level. It is possible that changes in *A* and *g_s_* seen at this sampling point are due to this drying effect and an increased rate of evapotranspiration under higher temperatures (Singh et al., 2015). Both *A* and *g_s_* rates in the ElevT plants returned to AmbT levels at 90d and 120d when both treatment groups were sufficiently watered, likely attributing the negative effects seen at 60d to water deficit.

**Figure 2.**
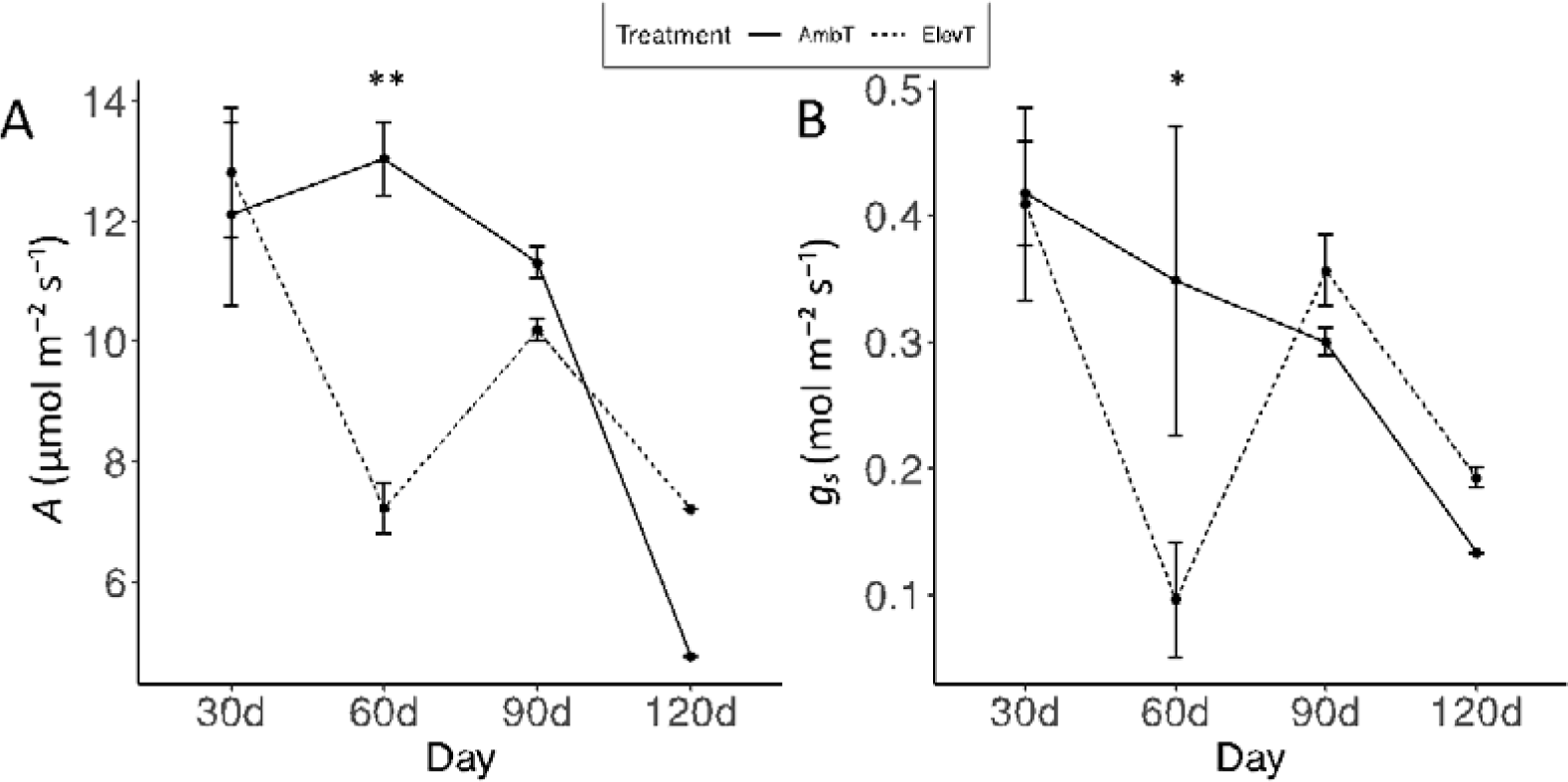
Gas exchange measurements of potato plants grown in AmbT and ElevT conditions. Rates of A) carbon assimilation (*A*, measured in umol m^-2^ s^-1^) and B) stomatal conductance (*g_s_*, measured in mol m^-2^ s^-1^) of potato plants are shown. Solid lines represent AmbT plants while dashed lines represent ElevT. Error bars represent standard error between biological replicates (*n* = 2). Asterisks indicate significant differences between AmbT and ElevT from pairwise *T*-tests at each time point (* = *p* < 0.05; ** = *p* < 0.01).

Chlorophyll fluorescence was also measured at each time point with the maximum PSII quantum efficiency (F_v_/F_m_) used as a proxy for rates of dark-induced leaf senescence (Zwack et al., 2016). There were no significant differences in senescence rates between treatment groups except at 60d, where the ElevT plants had 44.7% higher Fv/Fm measurements than AmbT leaves after 72hr (Supplementary Figure 2). This indicates a possible slower rate of senescence in the ElevT plants at 60d, which is the same time point when gas exchange measurements showed a significant decline in ElevT plants, likely due to the mild drought stress they were receiving.

Additionally, there were no significant differences in total leaf chlorophyll content between plants grown in AmbT and ElevT at any time points measured (Supplementary Figure 3).

### Biomass and Yield

At each collection time point, tubers were harvested from all chambers and assigned a size class based on weight: tuber initials (TI), immature tubers (IMT) and mature tubers (MAT) (Supplementary Fig. 1). TI were collected at each time point in both AmbT and ElevT conditions, while IMT and MAT were not found at the 30d collection point in either treatment (Figure 3). There were no significant differences in the total number of TI or IMT collected between AmbT and ElevT treatments across developmental time points (Supplemental Table 4). At 90d, no MAT from ElevT plants were collected and at 120d there were significantly fewer average number of MAT per plant in ElevT (2.63 ± 1.8) compared to AmbT (6.12 ± 1.1) (Supplementary Table 4).

**Figure 3.**
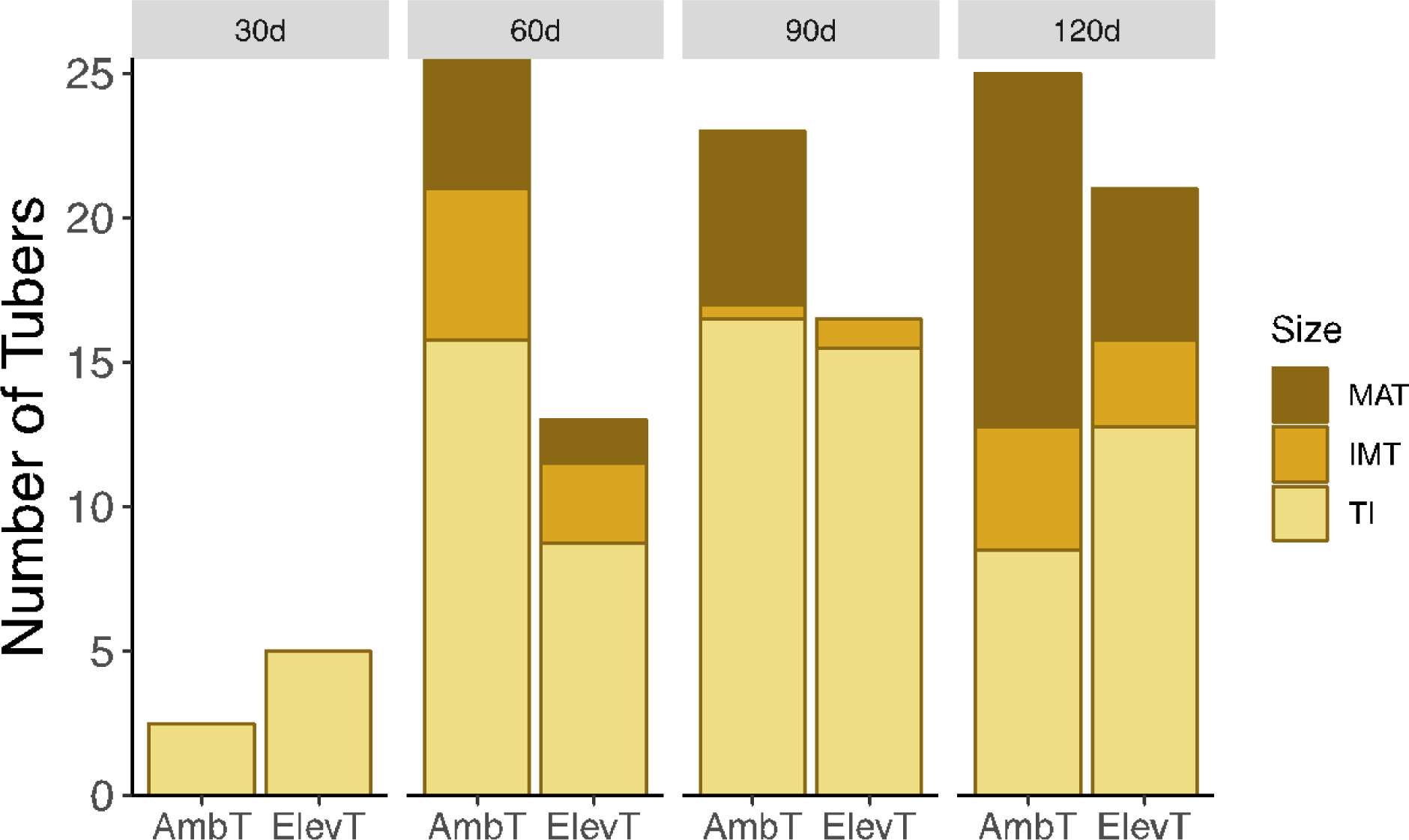
Stacked bar plot showing the average number of tubers per plant per treatment for each size class. Counts are shown as average number of tubers per treatment per size class. Two plants were harvested per chamber at 30 and 90d, and four plants per chamber were harvested at 60 and 120d, respectively.

At final harvest (120d), the average number of tubers per plant and average tuber weight per plant were determined for all tubers above 0.6 g (combining both IMT and MAT). Plants grown in ElevT conditions had significantly decreased tuber yield (both count and weight) compared to AmbT plants (Figure 4A-B). For tuber count, an average of 12.25 MAT per plant were collected from AmbT plants while in ElevT a significant decrease to 5.24 MAT per plant (57% decrease; Fig. 4A) was observed. Regarding tuber weight, AmbT plants had a mean tuber weight per plant of 210g ± 73 g compared to 72.3g ± 13.5 g for ElevT plants (66% decrease in ElevT; Fig. 4B). Plant height, aboveground FW, and aboveground DW were not significantly different between treatment groups, although the aboveground biomass of ElevT plants was slightly more than AmbT plants by up to 8% in both fresh and dry weight (Figure 4C-E).

**Figure 4.**
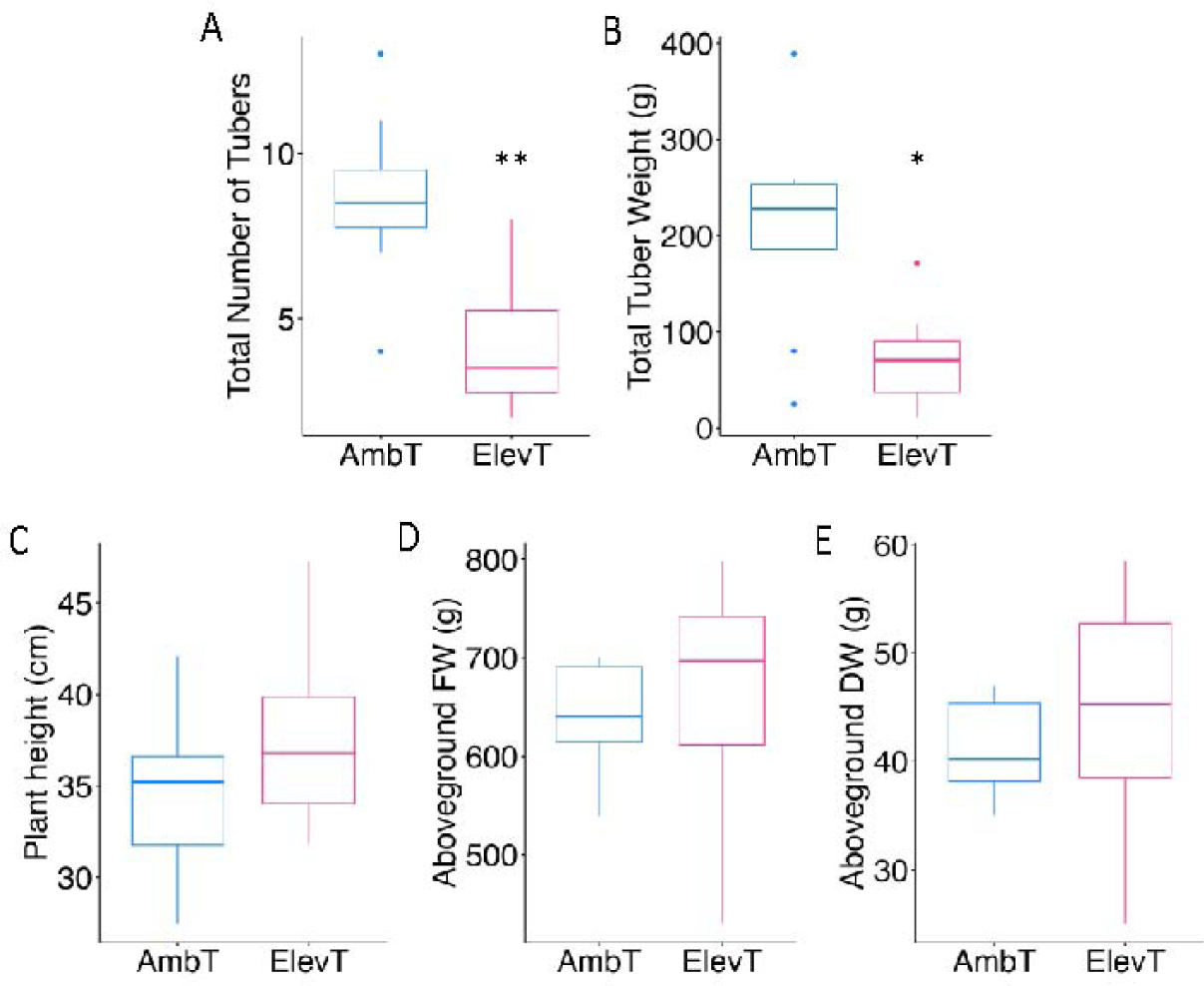
Final harvest data from potato plants grown in AmbT and ElevT conditions. Box plots showing the A) average total number of tubers per plant, B) average total tuber weight per plant, C) average plant height (cm), D) average aboveground fresh weight (FW) per plant (g), and E) average aboveground dry weight (DW) per plant (g) for each treatment group at 120d. Values represent the average of four plants per chamber (*n* = 2). Tuber initials were excluded from final yield measurements. Asterisks indicate significant difference from AmbT (* = *p* < 0.05; ** = *p* < 0.01).

### Phytohormone Content

To investigate changes in signaling in plants grown under ElevT, we analyzed endogenous phytohormone content of all tissues (leaf, TI, IMT, MAT) using LC/MS averaged across the 90d and 120d time points. In total, 49 phytohormone metabolite forms could be detected: 10 auxins, 23 cytokinins, 5 abscisic acids, 6 jasmonic acids (JAs), and 5 other phenolic compounds including salicylic acid (SA) (Supplementary Table 5A-D), with gibberellic acid forms below detectable levels. There were no significant differences in total levels of cytokinins, JAs, or SA between treatment groups, likely due to large variation among biological replicates (Supplemental Table 5A-D). Total auxin levels were significantly decreased in leaves, with a 46% decrease in leaves grown in ElevT conditions compared to AmbT conditions (Figure 5A). When specific auxin metabolites were compared, the total difference was likely attributable to significantly lower levels of oxo-indole-3-acetic acid (OxIAA) in leaves (55% decreased from AmbT to ElevT) (Fig. 5B).

**Figure 5.**
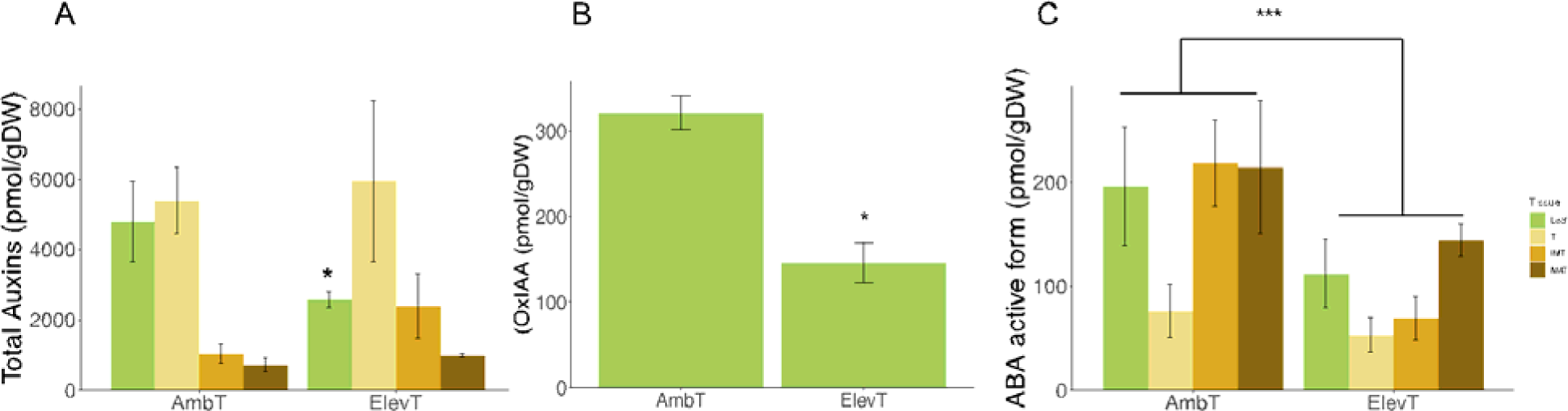
Total levels hormone levels in various tissues in potato in AmbT and ElevT conditions. A) Total levels of total auxins (pmol/gDW) across tissues; B) Levels of oxo-indole- 3-acetic acid (OxIAA) (pmol/gDW) in leaves; and C) Levels of the active form of ABA active across tissues pooled from 90 and 120d old plants. Asterisks indicate a significant difference from the corresponding tissue under AmbT (* = *p* < 0.05; *** = *p* < 0.0001). Error bars are standard errors between biological replicates (*n* = 2).

Due to its known role as both a stress hormone and its involvement in promoting tuber induction, the levels of ABA and its metabolites (ABAs) were also analyzed. The levels of ABA were significantly lower in ElevT compared to AmbT when averaged across all tissue types, with levels ranging between 76 to 219 pmol/g DW in AmbT and 53 to 144 pmol/g DW in ElevT (Figure 5C). The decrease in ABA levels is especially evident in IMT in which ElevT samples had a 68% decrease of ABA compared to AmbT (Fig. 5C).

### Differential Gene Expression Analysis

To examine differential expression of genes across the potato transcriptome, RNA-Sequencing (RNA-Seq) was performed on total RNA extracted from all tissue samples. Libraries were aligned to the tetraploid Atlantic Cultivar (ATL_V3) *S. tuberosum* genome and mapping alignment statistics were assessed (Supplementary Table 2). Principal component analysis (PCA) showed that most of the variation in RNA-seq libraries was accounted for by organ type, with differences between leaves and tubers explaining 91% of the variance (Supplementary Figure 4). To identify effects of each treatment on gene expression, a separate PCA was conducted for the samples within each treatment group. From the PCA of AmbT samples, distinct clusters of each tissue type were observed along with a clear developmental gradient of tuber sizes (Figure 6A). This was not observed however, in the PCA of ElevT, where the clusters of IMT and MAT are more overlapping (Figure 6B). This suggests that ElevT has a differential impact on development and tuber identity compared to AmbT with regards to gene expression.

**Figure 6.**
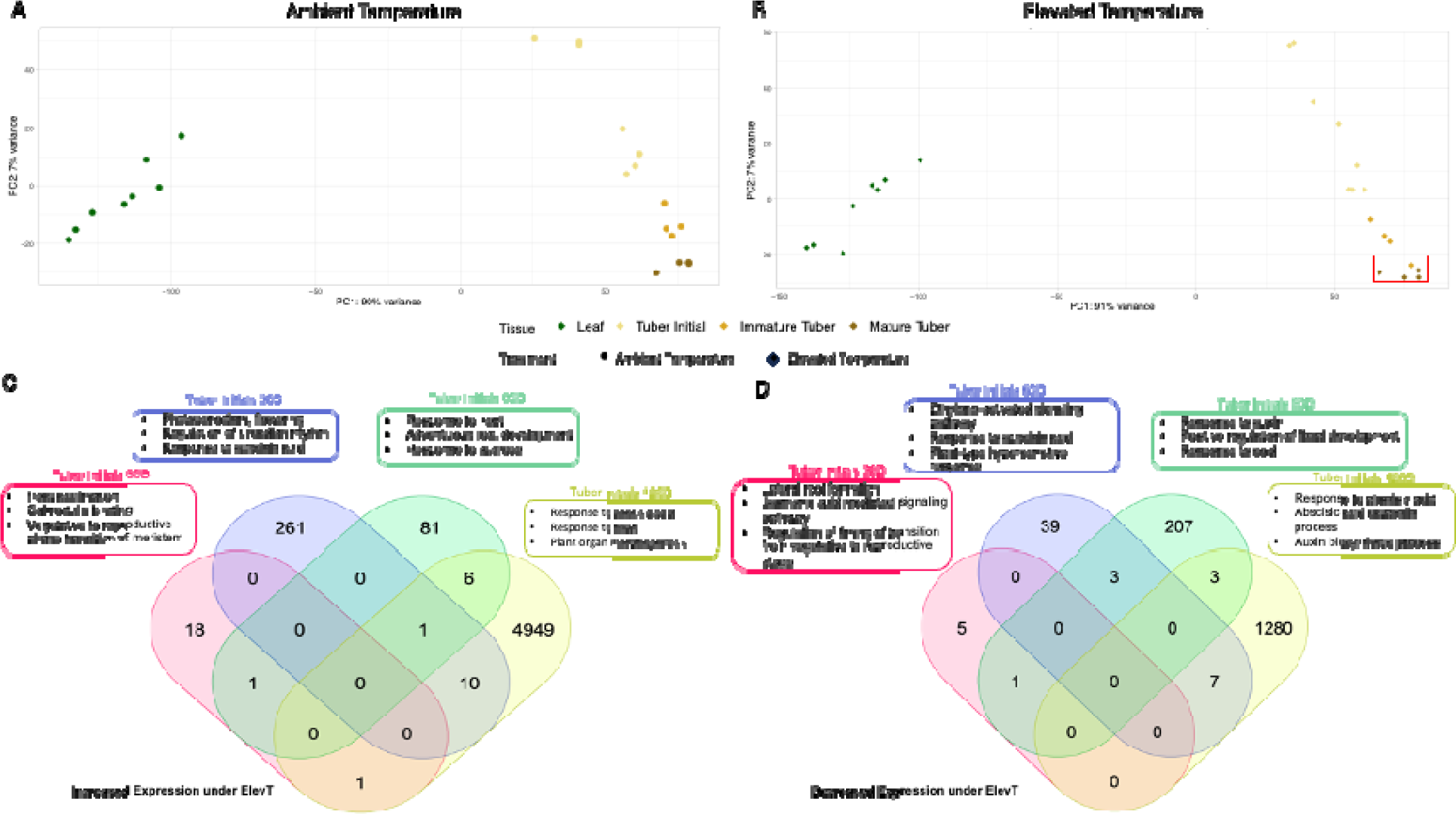
**PCA and venn diagram of expression patterns across RNA-Sequencing libraries**. PCA with libraries colored by tissue type for AmbT (A) and ElevT (B). Graphs were produced using *prcomp* in R. (C) Venn diagram highlighting the number of shared increased DEGs between developmental stages of tubers under both AmbT and ElevT, with selected GO terms of DEGs at each timepoint. (D) Venn diagram highlighting the number of shared decreased DEGs between developmental stages of tubers under both AmbT and ElevT, with selected GO terms of DEGs at each timepoint. Venn diagram was made using https://molbiotools.com/listcompare.php.

Differential expression analysis was then completed to determine genes with significant increases or decreases in expression across treatments, tissues and time points using *DESeq2* (Love et al. 2014). Genes with *p-*adjusted values of < 0.05 were considered differentially expressed. In leaves, there were a total of 1,078 differentially expressed genes (DEG) between ElevT and AmbT across all time points, with 15, 615, 146 and 302 DEGs at 30d, 60d, 90d, and 120d comparisons, respectively (Supplementary Table 6A-D). In TI, there were a total of 26, 321, 302, and 6,257 DEGs in the 30d, 60d, 90d, and 120d comparisons, respectively (Supplementary Table 7A-D).

GO association of biological processes was completed on DEGs in elevated temperature for TI. These processes were identified for both DEGs with increased and decreased expression at each timepoint (Figure 6C-D). DEGs with increased expression in TI in ElevT were involved in response to heat, regulation of circadian rhythm and calmodulin binding (Table 1). DEGs with decreased expression in TI in ElevT were involved in regulation of flower development, lateral root formation and response to cold (Table 2). More so, a clear overlap between phytohormone and transcriptome expression was seen in processes involving ABA and Auxin as shown in Figure 6C-D.

**Table 1.** Subset of DEGs with increased expression found in TI and their associated GO term.

| Timepoint | GO Term | ATL Transcript ID | ATL Functional Annotation |
| --- | --- | --- | --- |
| 30d | Heat Acclimation | Soltu.Atl_v3.12_4G008690.1 | Co-chaperone GrpE family protein |
|  | Calmodulin binding | Soltu.Atl_v3.10_3G006070.1 | lipid transfer protein |
|  | Vegetative to reproductive phase transition of meristem | Soltu.Atl_v3.06_2G022450.2 | ubiquiting-conjugating enzyme |
| 60d | Response to Absciscic Acid | Soltu.Atl_v3.S043390.1 | Transmembrane amino acid transporter family protein |
|  | Regulation of circadian rythim | Soltu.Atl_v3.01_1G022500.4 | Homeodomain-like superfamily protein |
|  | Photoperiodism, flowering | Soltu.Atl_v3.04_3G013030.1 | gigantea protein (GI) |
| 90d | Response to Heat | Soltu.Atl_v3.12_4G003540.1 | related to AP2 |
|  | Adventious Development | Soltu.Atl_v3.09_1G015110.1 | cytochrome P450, family 83, subfamily B, polypeptide |
|  | Response to sucrose | Soltu.Atl_v3.S027640.4 | alcohol dehydrogenase |
| 120d | Response to Absciscic Acid | Soltu.Atl_v3.09_4G023670.3 | Mitochondrial transcription termination factor family protein |
|  | Response to heat | Soltu.Atl_v3.12_2G026260.1 | temperature-induced lipocalin |
|  | Plant organ morphonogenesis | Soltu.Atl_v3.S105180.1 | AINTEGUMENTA-like |

**Table 2.**
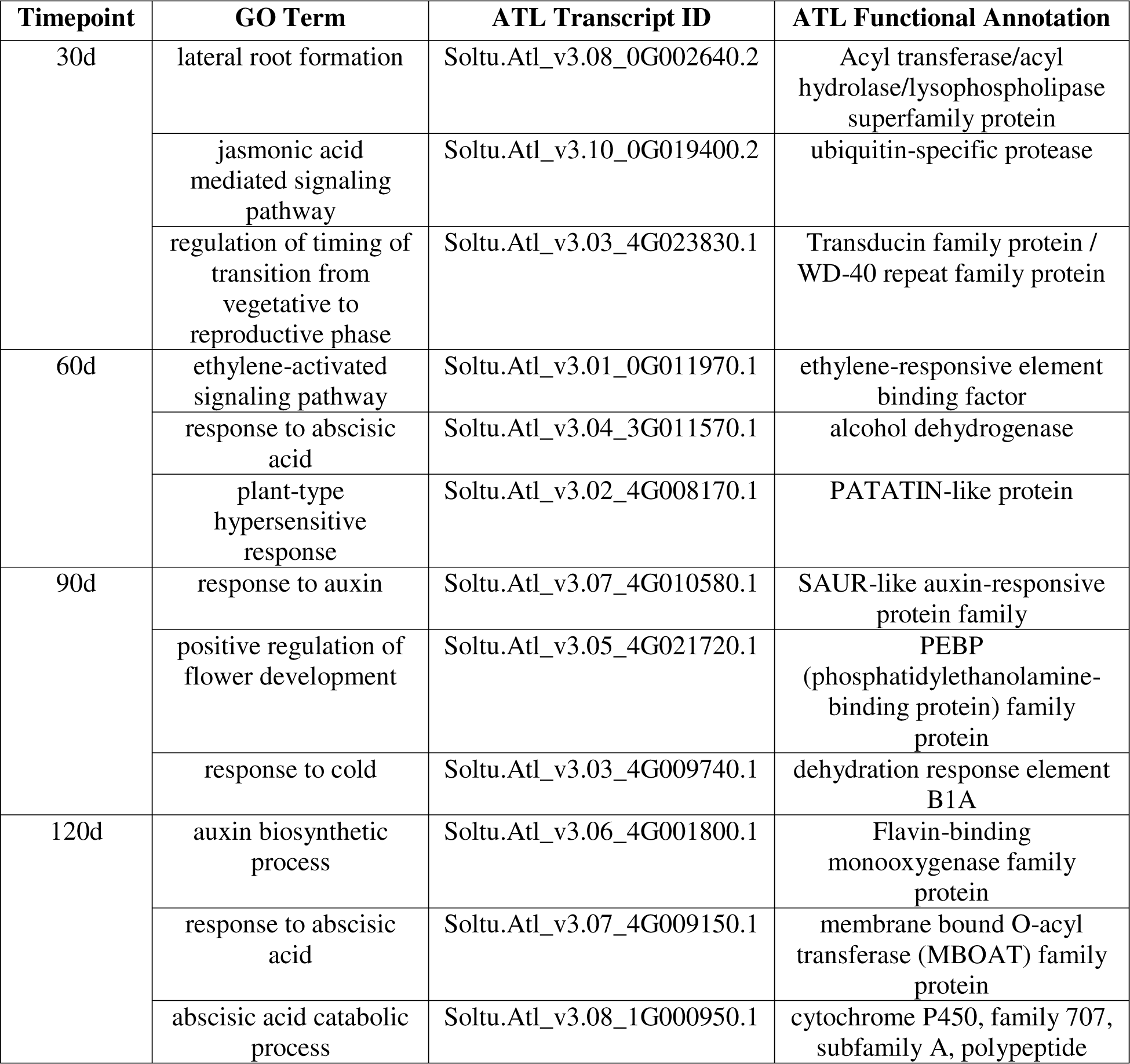
Subset of DEGs with decreased expression found in TI and their associated GO term.

| Timepoint | GO Term | ATL Transcript ID | ATL Functional Annotation |
| --- | --- | --- | --- |
| 30d | lateral root formation | Soltu.Atl_v3.08_0G002640.2 | Acyl transferase/acyl hydrolase/lysophospholipase superfamily protein |
|  | jasmonic acid mediated signaling pathway | Soltu.Atl_v3.10_0G019400.2 | ubiquitin-specific protease |
|  | regulation of timing of transition from vegetative to reproductive phase | Soltu.Atl_v3.03_4G023830.1 | Transducin family protein / WD-40 repeat family protein |
| 60d | ethylene-activated signaling pathway | Soltu.Atl_v3.01_0G011970.1 | ethylene-responsive element binding factor |
|  | response to abscisic acid | Soltu.Atl_v3.04_3G011570.1 | alcohol dehydrogenase |
|  | plant-type hypersensitive response | Soltu.Atl_v3.02_4G008170.1 | PATATIN-like protein |
| 90d | response to auxin | Soltu.Atl_v3.07_4G010580.1 | SAUR-like auxin-responsive protein family |
|  | positive regulation of flower development | Soltu.Atl_v3.05_4G021720.1 | PEBP (phosphatidylethanolamine-binding protein) family protein |
|  | response to cold | Soltu.Atl_v3.03_4G009740.1 | dehydration response element B1A |
| 120d | auxin biosynthetic process | Soltu.Atl_v3.06_4G001800.1 | Flavin-binding monooxygenase family protein |
|  | response to abscisic acid | Soltu.Atl_v3.07_4G009150.1 | membrane bound O-acyl transferase (MBOAT) family protein |
|  | abscisic acid catabolic process | Soltu.Atl_v3.08_1G000950.1 | cytochrome P450, family 707, subfamily A, polypeptide |

### Expression Patterns of Known Regulators in Tuberization Signaling

We then investigated the gene expression patterns of key genes and known regulators involved in tuberization signaling in potato. For each key gene of interest, we provide expression information for one syntelog in this section. Syntelogs are homologous genes resulting from a whole-genome duplication event or speciation and are expected to share similar functions.

Supplementary Figure 5A-G provides expression data for all syntelogs associated with each gene of interest and individual results from our syntelog analysis are in Supplementary Table 3. The genes of interest observed that induce tuberization include *StSP6A* (Soltu.Atl_v3.05_1G024440.1), *StBEL5* (Soltu.Atl_v3.06_3G019920.1) and *StIT1* (Soltu.Atl_v3.06_2G021580.1) (Figure 7A-C and Supplementary Figure 5A-C). Expression of key genes known to inhibit tuberization were also analyzed, including *StCOL1* (Soltu.Atl_v3.02_1G027420.1), *StSP5G-A/B* (Soltu.Atl_v3.05_3G022530.1), and *StTOC1* (Soltu.Atl_v3.06_4G015370.2) (Figure 7D-F and Supplementary Figure 5D-G).

**Figure 7.**
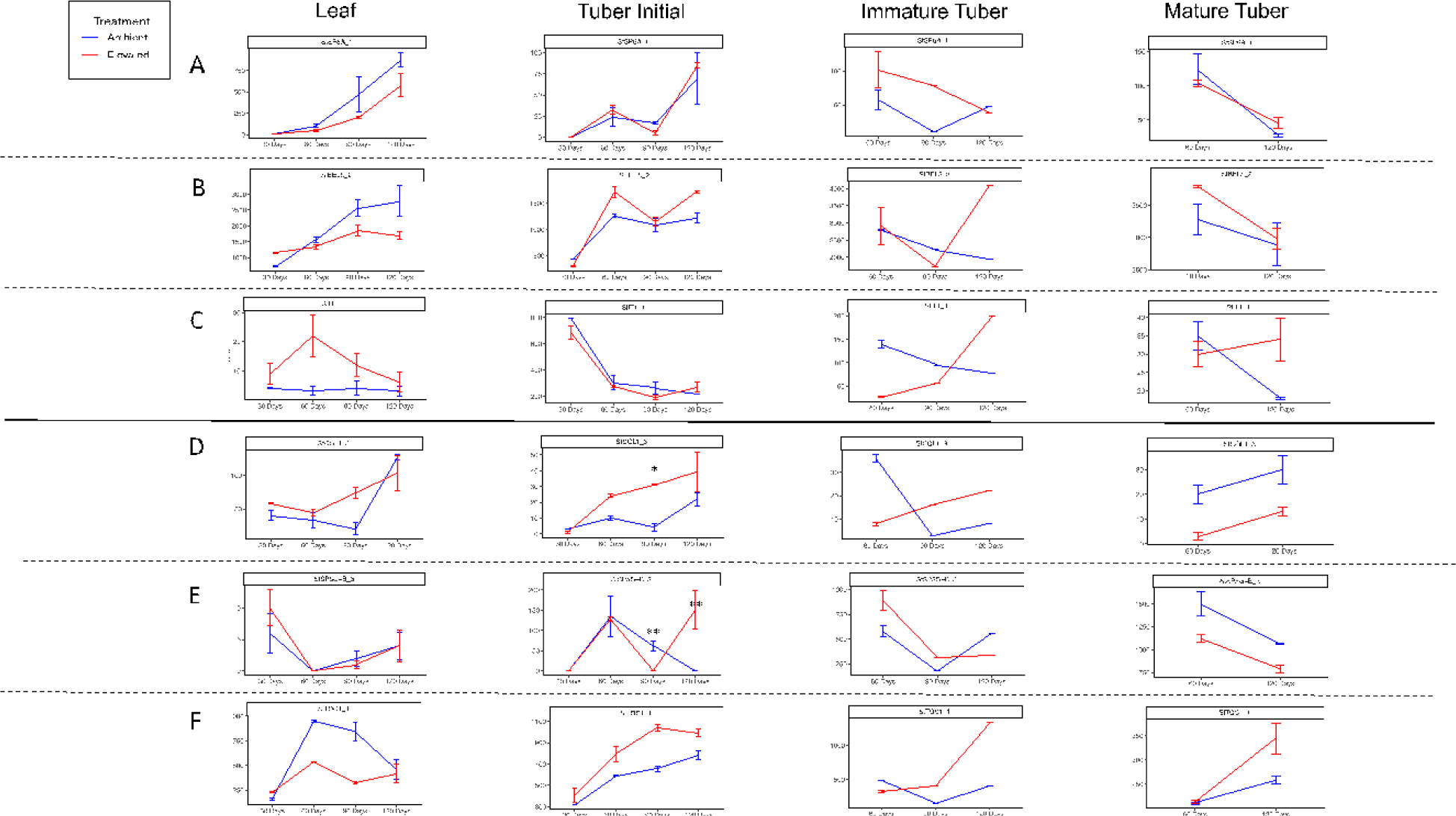
Transcript abundance (TPM) of known tuberization genes in each tissue type. A- C: Tuberization promoting genes (A - *StSP6A*, B- *StBEL5*, and C - *StIT1*). D-F: Tuberization inhibitor genes (D-F: D- *StCOL1*, E- *StSP5G-B*, and F-*StTOC1,).* TPM values are presented each tissue type for each timepoint over the course of 120 days. Solid lines represent AmbT plants while dashed lines represent ElevT. No ElevT mature tubers were collected at 90d, so no data is shown for that point. (* = *p* < 0.10; ** = *p* < 0.05).

*StSP6A* expression in leaves was consistent with its known gradual expression pattern (Park et al., 2022) amongst syntelogs and tissues in both treatments but was not differentially expressed in the ElevT treatment (Figure 7A & Supplementary Figure 5A). *StBEL5*, which is known to be a transcriptional activator that targets *StSP6A* was also not differentially expressed at any timepoint (Sharma et al., 2016). Additionally, the expression pattern of *StBel5* syntelogs were observed to be almost identical with exception of syntelog *StBEL5_3* (Fig. 7B & Supplementary Figure 5B).

In AmbT, *StIT1* expression was highest in TI at 30d, then decreases by 61% by 60d and eventually decreases to its lowest expression in AmbT at 120 time point (Fig. 7C). Due to the early role of *StIT1* in promoting tuberization of stolons, we would expect expression of this gene to be lower at later stages of tuber development (IMT and MAT), which is what was observed (Fig. 7C). *StIT1* had relatively lower abundance in ElevT in TI tissue but was not identified as differentially expressed due to ElevT.

In terms of repressors of tuber formation, one *StCOL1* syntelog was observed to have increased expression at 90d in TI (Figure 7; *p* < 0.08). Additionally, some syntelogs of *StSP5G-B* in TI were differentially expressed at both 90d and 120d, while all syntelogs were observed to have significant increase in expression at 120d in ElevT. *StSP5G-B* has been previously identified as a downstream target of *StCOL1*, and to be differentially expressed by elevated temperature (Hancock et al., 2014). *StTOC1* showed increased expression in all tuber classes under ElevT but was not significantly differentially expressed (Fig 7F). In leaves, we saw non-significant reduction in expression of *StTOC1* at all timepoints except at 30d (Fig. 7F).

## Discussion

### Tuber yield loss under heat stress is not caused by a decrease in photosynthetic parameters

In this study, we investigated the mechanisms of yield loss that is observed when potato plants are exposed to chronic elevated temperature. From leaf physiology measurements we found that *A* and *g_s_* were not significantly decreased in ElevT compared to AmbT at every time point other than 60d when these plants were found to have drier soil than in AmbT (Fig. 2). Upon experiencing water deficit, plants produce signals to close their stomatal pores, thus reducing transpiration and conserving water, which significantly reduces gas exchange of leaves (Galmés et al., 2013). As such, the significantly lowered rates of *A* and *g_s_* at the 60d time point are most likely due to insufficient watering. Our results also suggest that future project elevated temperatures for potato do not significantly impact leaf senescence (as seen through changes in F_v_/F_m_) or chlorophyll content when water availability is not a concern (Supplementary Figs. 2, 3). These results are consistent with other studies that found no significant or positive effects of high temperatures on photosynthetic rates and chlorophyll content in potato (Hancock et al., 2014; Park et al., 2022).

Despite lack of significant effects of elevated temperature on leaf physiology in our experiment, we still observed a significant decrease in tuber number and yield from ElevT plants (Fig. 3; Fig. 4; Supplementary Table 4). This implies that elevated temperatures affect the aboveground and belowground components of potato plants differently. Moreover, many studies indicate significant increases in aboveground biomass of potato plants exposed to high temperatures (Tang et al., 2018; Timlin et al., 2006). Although there were no significant differences in aboveground biomass observed in this study, measurements increased slightly under ElevT (Fig. 4C-E), consistent with findings in literature. Thus, results from our study suggest a reduction in photoassimilate production is not a driver of yield loss under heat stress.

### Changes in phytohormone content in potato under heat stress may impact tuberization

Internal signaling in potato plants is crucial for the formation of tubers and maintenance of sink strength, both through genetic regulation and through hormone signaling (Dutt et al., 2017). In the present study, our phytohormone analyses revealed several changes in phytohormone content. This included significantly lower levels of ABA active form across all tissue types under ElevT (Fig. 5C). Besides its role in stress response, ABA contributes to tuber development in *S. tuberosum* through antagonistic action on GA (Chen et al., 2022; Li et al., 2021). GA is an important inhibitor of tuberization through inducing stolon elongation which in turn inhibits tuber induction by repressing radial growth of stolons into tubers (Xu et al., 1998). ABA is also known to regulate dormancy of potato tubers, with lower levels generally associated with shorter dormancy periods (Wang et al., 2020). In this regard, the decreased ABA levels observed in tubers under elevated temperatures could explain the reports of heat sprouts and early dormancy breaks seen in potatoes grown under high temperature in other studies (Zhang et al., 2021).

Recent work showed that StABI5-like 1(*StABL1*), a transcription factor central to ABA signaling, is a binding partner of FT-like genes *StSP6A* and *StSP3D*. When interacting with StSP6A and StSP3D, it was shown that an alternative tuberigen activation complex is formed and promotes tuberization (Jing et al., 2022). Overexpression of this *StABL1* gene resulted in earlier flowering, tuberization and overall shorter plant life cycle. Coupled with the finding of ABA-associated DEGs from our TI tissue (Fig. 6C-D), these results indicate that further work is needed to understand the role of ABA signaling in tuber development under higher temperatures.

### Chronic elevated temperatures inhibit tuber filling, but not initiation

Tuber production of a potato plant consists of three main separate processes: tuber formation/initiation, tuber filling, and tuber maturation stage (Liu et al., 2020). Quantification of the number of TI and IMT produced under ElevT showed no significant change compared to AmbT (Fig. 3; Supplementary Table 4). However, we observed an increased average number of TI at 30d in ElevT compared to AmbT (Fig. 3). These results suggest that tuber initiation is not significantly inhibited under chronic elevated temperatures. Nevertheless, we did observe a significant decrease in MAT yield collected from ElevT plants (Fig. 3; Supplementary Table 4). This implies a disruption in the normal developmental process under ElevT, hindering tubers from reaching maturity or filling adequately. Past studies with acute ElevT treatments have shown it negatively impacts potato yield by reducing amount of fully matured potato tubers (Kim & Lee, 2019).

Additionally, there were observable differences between RNA-seq clustering between treatments. Plants grown in ElevT conditions showed stronger clustering of MAT and IMT libraries compared to MAT and IMT samples from plants grown in AmbT conditions (Figure 6 A-B). This indicates that IMT from plants grown in ElevT conditions exhibit more similar transcript expression patterns to MAT than when grown in AmbT. This aligns with previous reports that identified an increase in physiological age of tubers because of elevated temperatures (Wiersema, 1985). Therefore, chronic heat stress increases the developmental age of tubers which in turn decreases the potato in-season growing time and its post-harvest tuber dormancy time (Wiersema, 1985). This implies that heat stress caused by future climate change likely disrupts normal signaling patterns that occur between tuber initiation and maturation, impacting tuber development and ultimately yield.

### Changes in expression of tuberization genes were observed in elevated temperature

There has been considerable work in recent years on understanding the effects of elevated temperature on the tuberigen *StSP6A* expression and its role in tuberization (Lehretz et al., 2017; Park et al., 2022). A previous study observing transgenic *StSP6A* over-expressor lines concluded that *StSP6A* likely controls tuber formation, but not continued growth of tubers as yield was still significantly decreased in transgenic *StSP6A* over-expressor lines under high temperatures (Park et al. 2022). In this study, there were no significant differences with expression of *StSP6A* under ElevT albeit there was a consistent decrease in *StSP6A* abundance in this treatment (Figure 7A). In the current study we found the tuberigen repressor, *StSP5G-A* had relatively higher expression in leaves at 30d and had significantly increased expression under ElevT at 90d in TI (Fig. 7E). In our results, we saw preferential expression of *StSP5-A* in leaves while *StSP5G-B* had higher expression in tuber classes under both treatments (Supplementary Fig. 5E-F). These results align with past studies that have observed higher expression of *StSP5G* under ElevT in leaves, while we also observed them to be DEGs under TI (Hancock et al., 2014).

Moreover, *StCOL1* which is known to induce expression of *StSP5G*, was observed to have increased expression in TI under ElevT at 90d (Fig. 7D). *StCOL1* is a CONSTANS-like gene part of a well described CONSTANS/FLOWERING LOCUS T (CO/FT) module that is central for sensing photoperiod and regulating developmental processes within Arabidopsis and more recently in other plants (Turck et al., 2008). From our data, in AmbT we see similar expression patterns between FT-like *StSP6A* and *StCOL1* in leaves, which is crucial for proper photoperiod sensing and leads to proper tuber development and increased tuber yield (Supp. Fig. 5A,5D; Fig. 3). On the other hand, expression of *StCOL1* in leaves grown in ElevT conditions shows a distinct pattern to *StSP6A* across sampled time points (Supp. Fig. 5A, 5D). The contrasting patterns of *CO/FT* are particularly crucial in leaves since they serve as the primary site for photoperception, where the CO/FT module has an effect in plant development (Valverde, 2011). We can then infer that divergence in the patterns between AmbT and ElevT in leaves is possibly leading to an incorrect perception of environmental cues necessary for proper tuber development in ElevT. Taken together, *StCOL1* may be involved both in misperception of environmental signals at ElevT through improper coordination of the *StCOL1/StSP6A* module in leaves as well as contributing to the reduction of tuber yield through induction of tuber repressors (*StS5PG*) in the same ElevT conditions.

One other inhibitor of *StSP6A* has been recently found to have increased expression in ElevT of *in-vitro* tuberization studies is *StTOC1* (Morris et al., 2019). From our RNA-seq data, we found no significant difference in expression of *StTOC1* albeit we observed its expression to increase over time and remain high in all tuber tissue classes under ElevT compared to AmbT (Fig. 7F). Within IMT and MAT we can also observe opposite expression patterns of *StTOC1* with *StSP6A*, indicating *StTOC1* may inhibit transcription of *StSP6A* in IMT and MAT near the end of the growing season. This data is consistent with previous studies that have found increases in expression of *StTOC1* in tubers under higher temperatures (Hancock et al. 2014). These results further highlight *StTOC1* as a potential gene responsible for heat tolerance in potato plants.

One tuberization pathway that remains unexplored under ElevT is through IDENTITY OF TUBER, *StIT1*. *StIT1* is a gene found to both interact with *StSP6A* and induce tuberization under non-inductive conditions (Tang et al., 2022). From our data, we observed non-significant difference in expression due to ElevT although we observed consistently less expression of *StIT1* in all tuber tissues under ElevT except at the 120d timepoint. Notably, it is at the 120d timepoint when we see a recovery of tuber yield, as at 90d we saw no MATs had developed in our ElevT treatment. Future work is needed to understand the possible role of *StIT1* in driving tuberization even under chronic elevated temperature conditions.

## Conclusion

The expected rise in global temperatures in coming decades poses an impending threat to the production of many staple crops, especially ones that are as susceptible to heat stress such as *S. tuberosum*. Understanding the effects of abiotic stress on crop plants is thus a crucial goal for coping with climate change. Here, we take a whole-plant approach to investigate the mechanisms of potato yield loss under heat stress using a realistic projection of future climate for major potato growing regions. Leaf-level physiology, endogenous phytohormone levels, and yield of potato plants grown under chronic ElevT were measured. Leaf gas exchange parameters (*A* and *g_s_*) were not significantly negatively impacted by ElevT, nor were leaf chlorophyll content or rates of leaf senescence. We observed a significant decrease in leaf ABA and auxin levels in leaves and tuber tissues under ElevT conditions, which indicates a possible role of these hormones in tuberization signaling in potato grown in ElevT. ElevT conditions lead to a reduction in the average number of tubers per plant at all timepoints except at 30d. From this we hypothesize that chronic ElevT is impacting tuber yield by inhibiting proper tuber filling and not by inhibiting tuber initiation. We also characterized expression of genes under chronic ElevT that are known contributors to tuberization. Results from this analysis contribute to the intricate understanding of the genetic regulation of tuberization under elevated temperatures in potato plants, highlighting the potential role of the *StCOL1-StSPGB* pathway as one target for generating heat tolerant potato cultivars (Figure 8). Other potential candidate genes to be explored are *StTOC1* and *StIT1,* offering different genetic pathways to sustain and enhance potato yield in the face of global warming and elevated temperatures.

**Figure 8.**
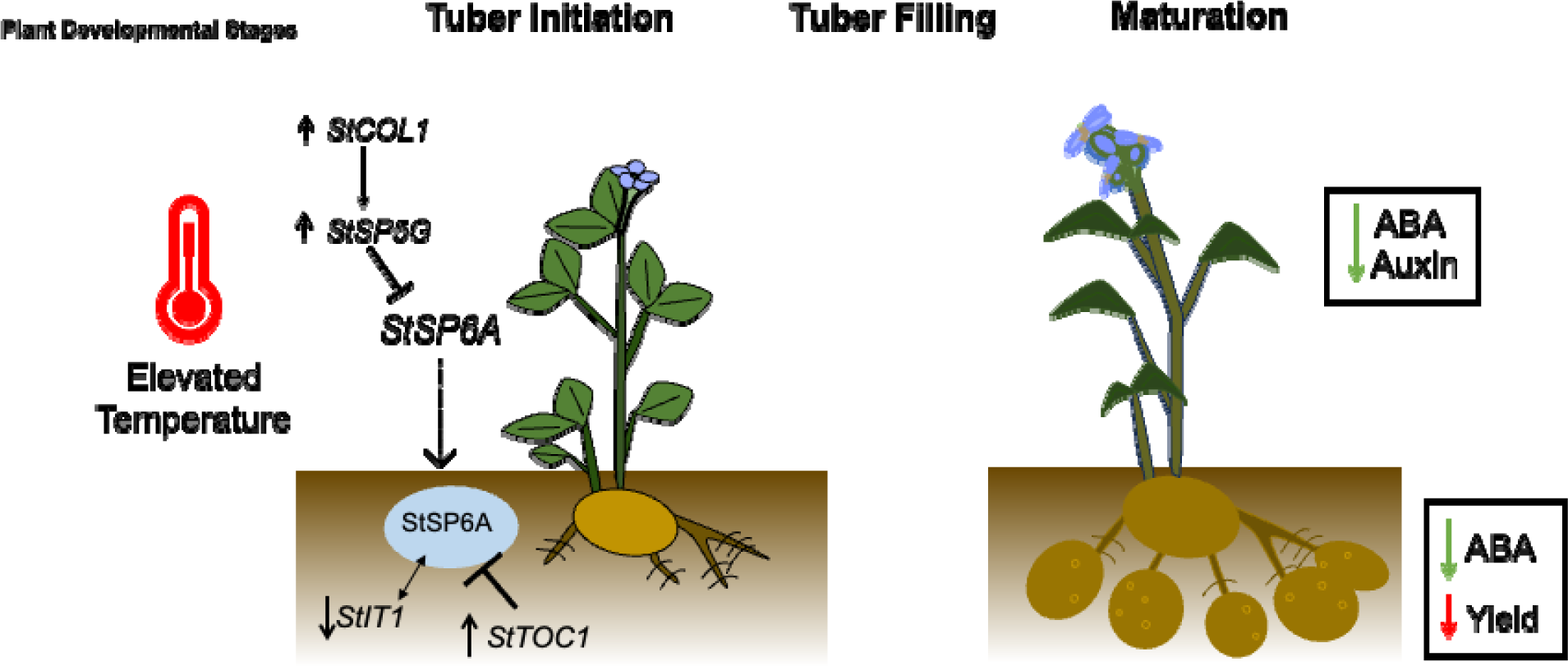
Model for known and potential changes in gene expression and hormone levels under elevated temperature. Dashed lines with arrows indicate movement, small arrows indicate increases or decreases in expression under elevated temperature. Lines with closed ends indicate inhibition while double arrowed lines between genes/proteins indicate interaction. Enclosed text are metabolite and plant physiology related parameters corresponding to above and belowground changes. Blue oval represents protein version of a *StSP6A*. *StSP6A*, FLOWERING LOCUS T homologue SELF PRUNING 6A; *StSP5G,* small RNA FT*-*homolog inhibitor; *StTOC1*, TIMING OF CAB EXPRESSION 1; StIT1, IDENTITY OF TUBER 1; ABA, Abscisic Acid.

## Acknowledgements and Funding

The authors would like to acknowledge USDA NIFA Hatch Project 1018601 to Courtney Leisner for supporting this work and the Ministry of Education, Youth and Sports of the Czech Republic from European Regional Development Fund-Project “TowArds Next GENeration Crops ” (CZ.02.01.01/00/22_008/0004581) for financing the hormonal analyses.

## Author Contributions

AMG, CPL and AMR designed the experiments. AMG collected physiology and RNA-Sequencing data. EP, PD, VM performed the phytohormone analysis. JPH generated the TAIR and GOSLIM annotations for the Atlantic reference genome. AMG, GHC, CPL analyzed and interpreted the data. AMG, GHC wrote the manuscript. All authors reviewed and approved the submitted version.

---

**Supplementary Figure 1.**
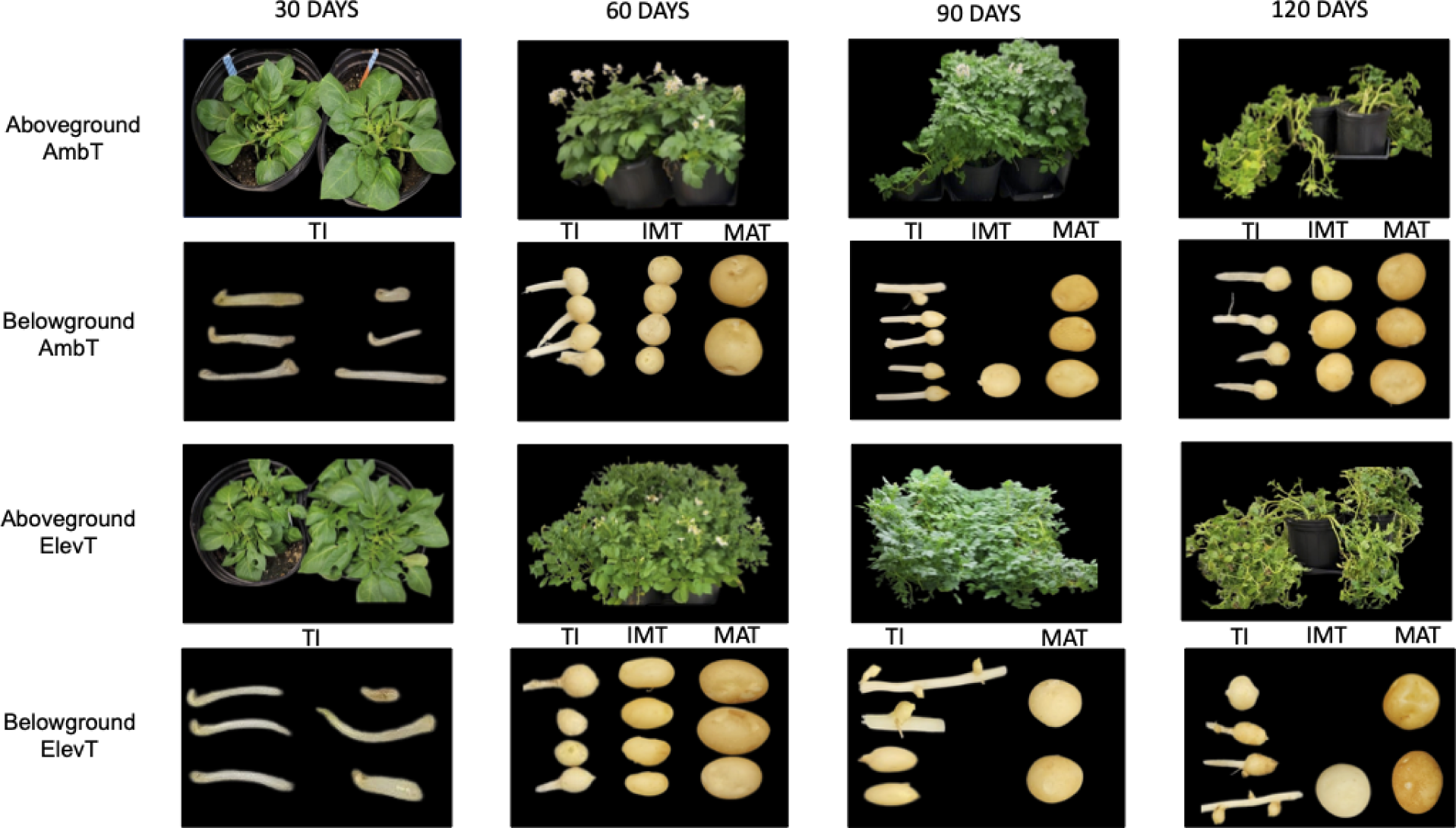
Example of tuber size classes for this experiment. Photographs of tuber initials (TI), immature tubers (IMT), and mature tubers (MAT) collected from AmbT and ElevT conditions throughout the growth chamber experiment.

**Supplementary Figure 2.**
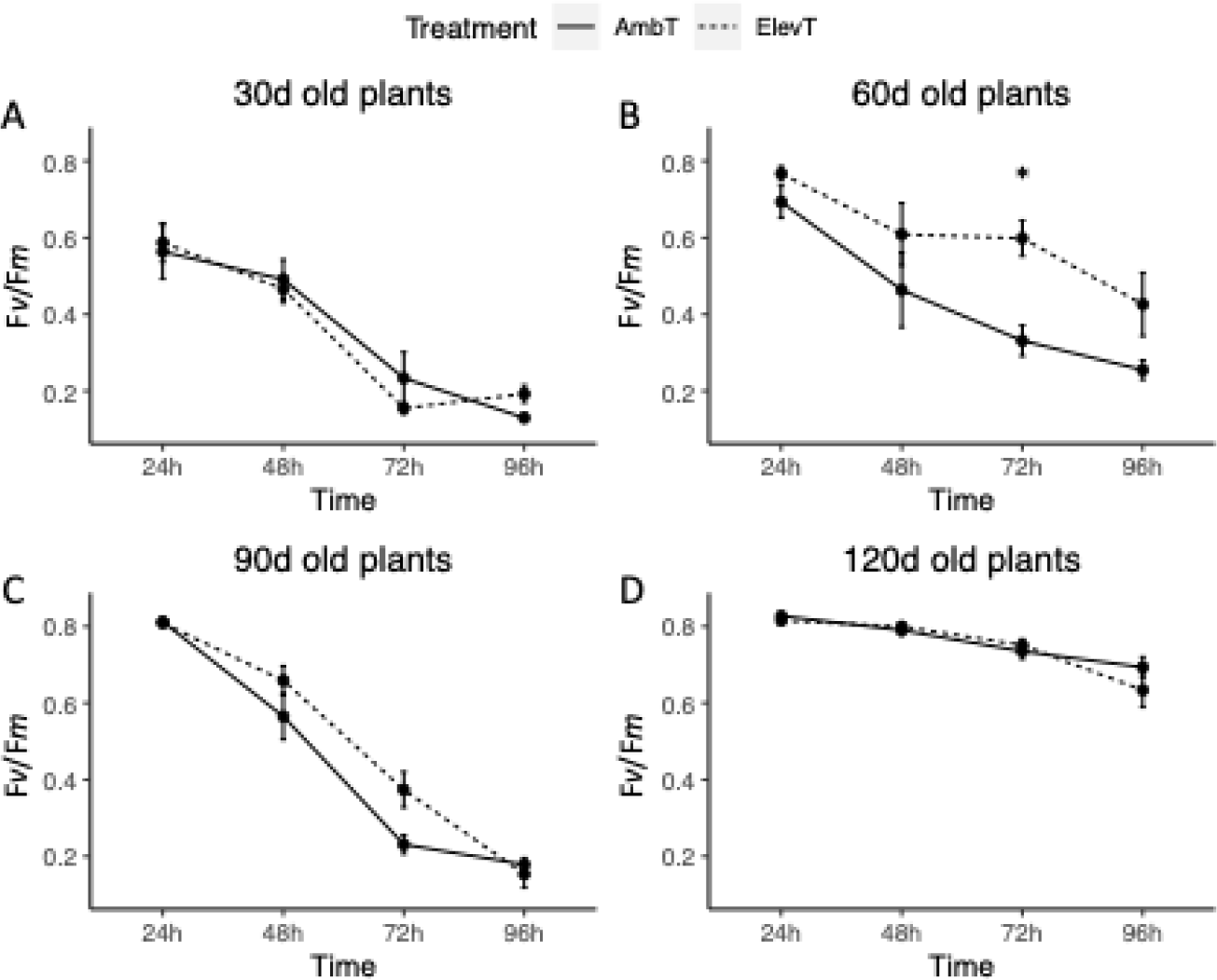
Measurement of maximum PSII efficiency (Fv/Fm) in AmbT and ElevT. Measurements were taken on leaves over the course of 96h to represent leaf senescence. Error bars are standard error between biological replicates (*n* = 2). Asterisks indicate significant differences between AmbT and ElevT from pairwise *T-*tests at each hour of measurement (** = *p* < 0.01).

**Supplementary Figure 3.**
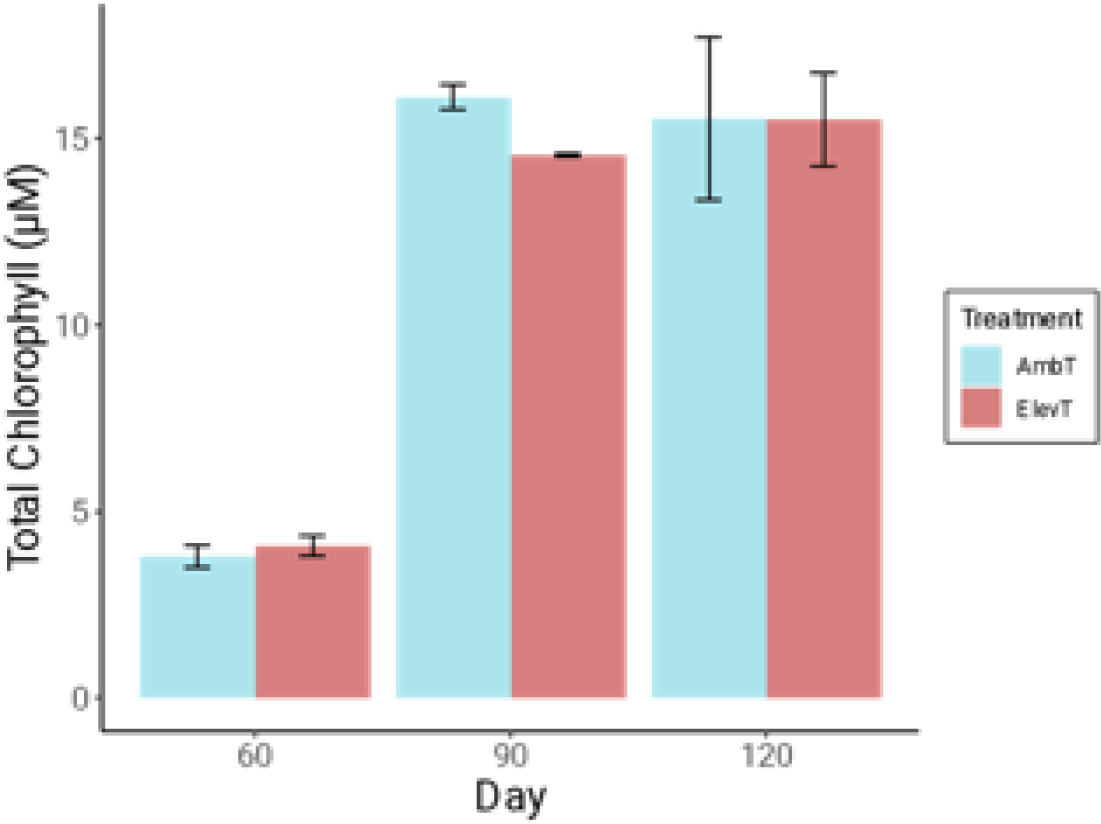
**Leaf total chlorophyll (**μ**M) content measured in AmbT and ElevT treatments.** Total chlorophyll values are averaged per four 6mm leaf discs collected from 60, 90, and 120d old plants. Error bars indicate standard error between biological replicates (*n* = 2). Pairwise *T*-tests were completed between treatments at each time point.

**Supplementary Figure 4.**
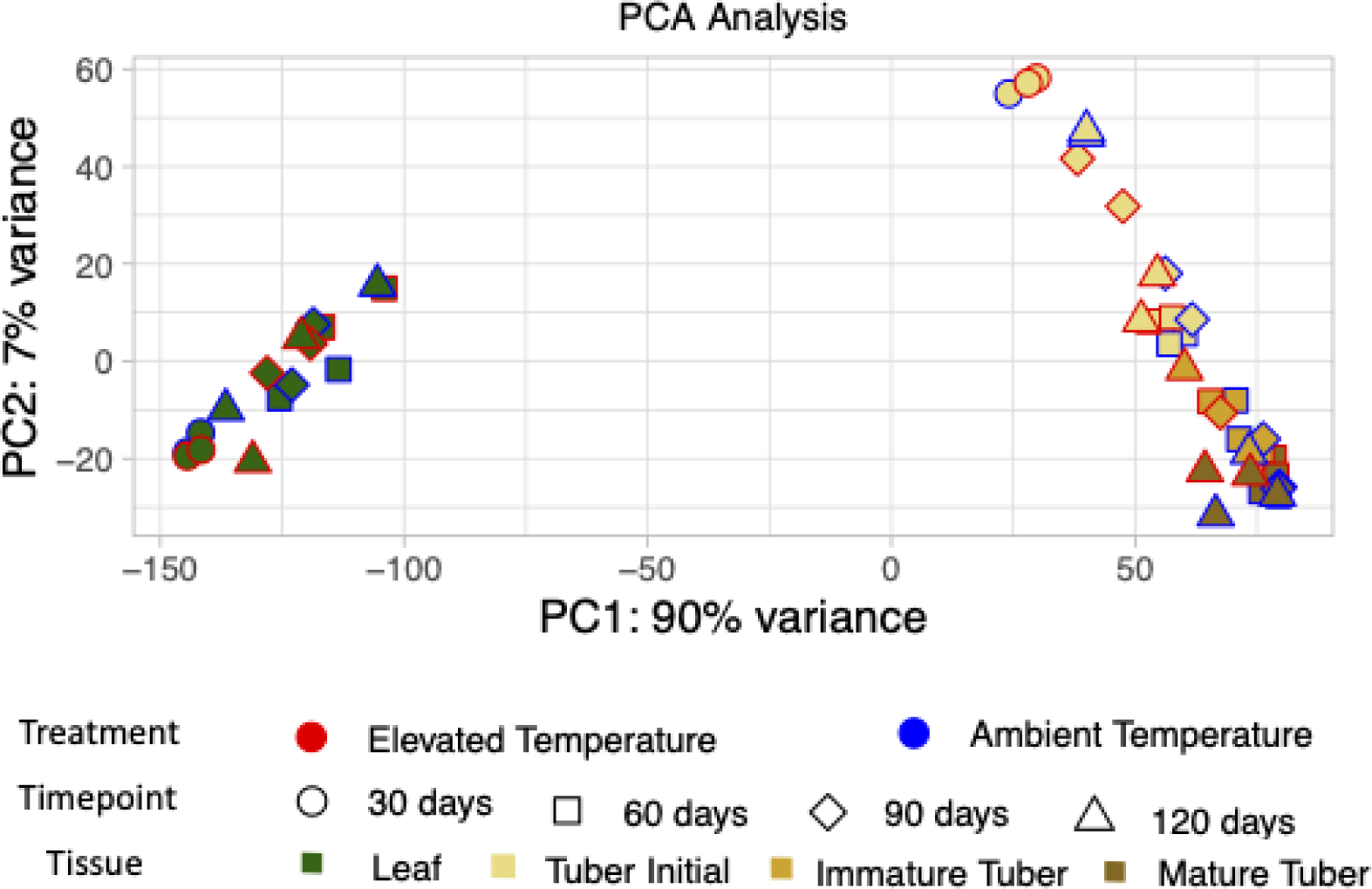
**Principal component analysis (PCA) of all RNA-Sequencing libraries**. Library samples are identified by treatment, timepoint and tissue. Graphs were produced using *ggplot* in R.

**Supplementary Figure 5A.**
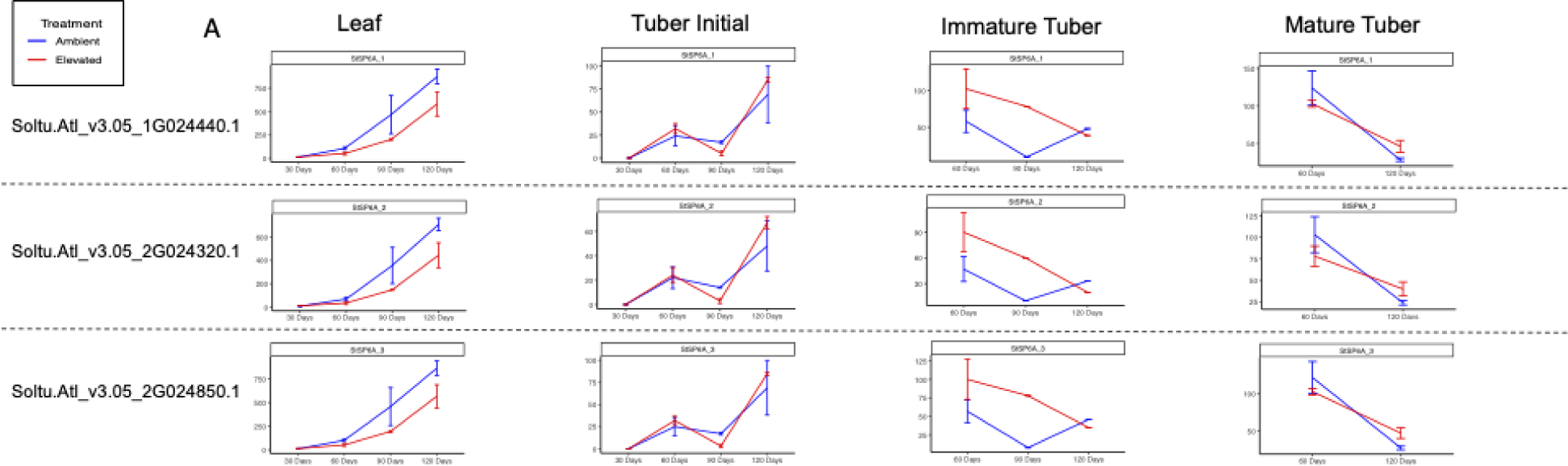
TPMs of known tuberization promoter gene *StSP6A* syntelogs in each tissue type over the course of 120 days. Solid lines represent AmbT plants while dashed lines represent ElevT. No ElevT mature tubers were collected at 90d, so no data is shown for that point. TPM values were determined from *Salmon* (Patro et al. 2017) and averaged per biological replicate (*n* = 1 or 2) (*p* < 0.10 = *; *p* < 0.05 = **).

**Supplementary Figure 5B.**
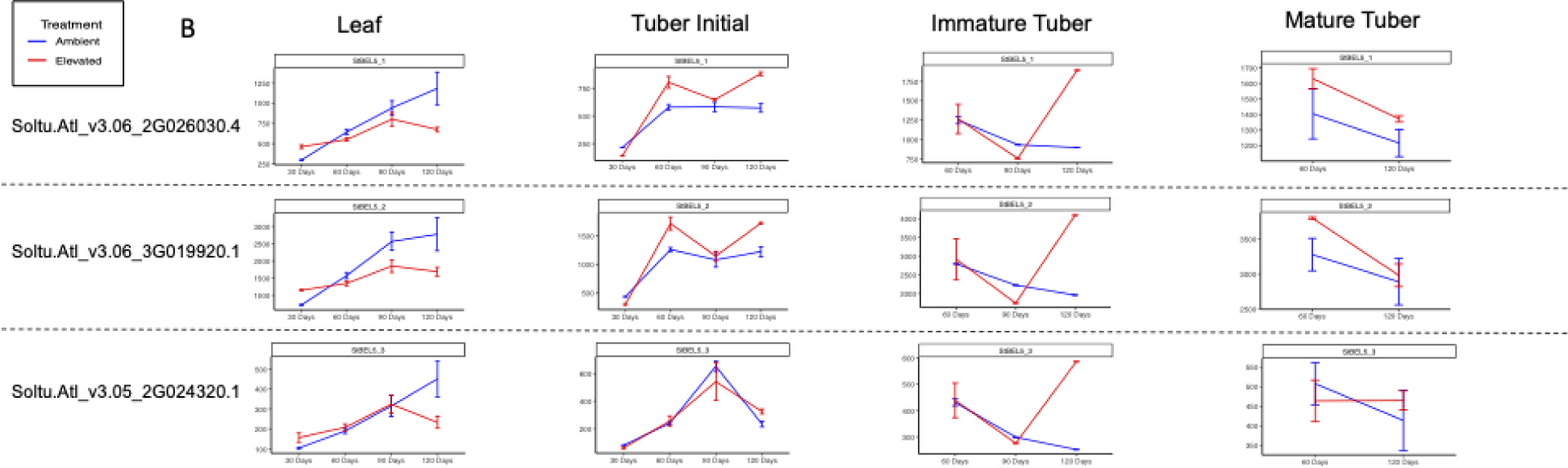
TPMs of known tuberization promoter gene *StBEL5* syntelogs in each tissue type over the course of 120 days. Solid lines represent AmbT plants while dashed lines represent ElevT. No ElevT mature tubers were collected at 90d, so no data is shown for that point. TPM values were determined from *Salmon* (Patro et al. 2017) and averaged per biological replicate (*n* = 1 or 2) (*p* < 0.10 = *; *p* < 0.05 = **).

**Supplementary Figure 5C.**
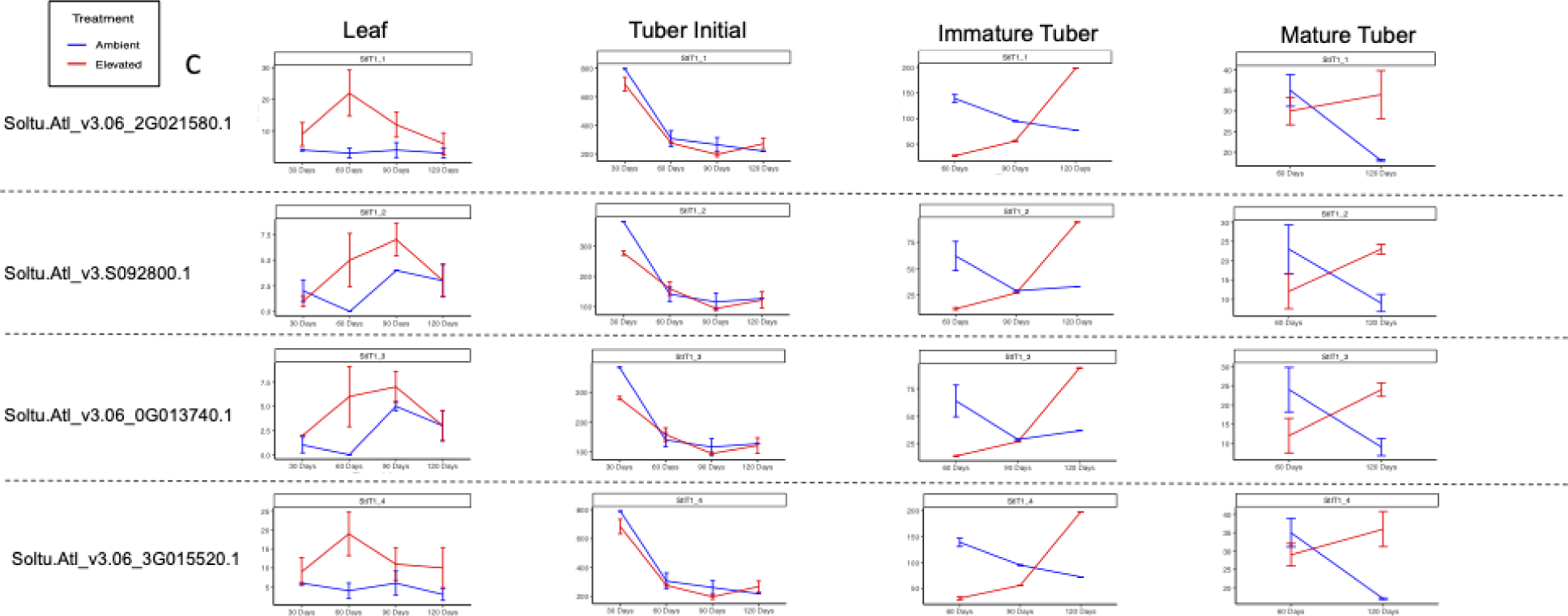
TPMs of known tuberization promoter gene *StIT1* syntelogs in each tissue type over the course of 120 days. Solid lines represent AmbT plants while dashed lines represent ElevT. No ElevT mature tubers were collected at 90d, so no data is shown for that point. TPM values were determined from *Salmon* (Patro et al. 2017) and averaged per biological replicate (*n* = 1 or 2) (*p* < 0.10 = *; *p* < 0.05 = **).

**Supplementary Figure 5D.**
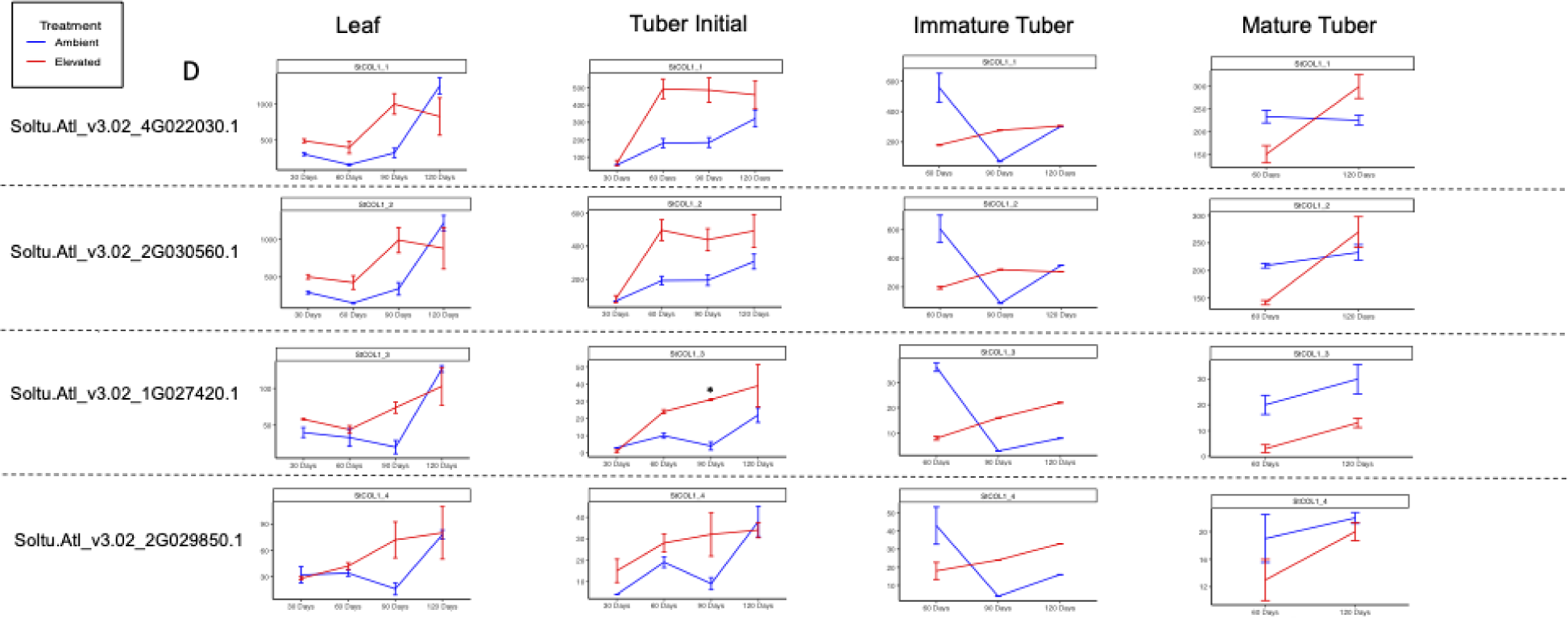
TPMs of known tuberization inhibitor gene *StCOL1* syntelogs in each tissue type over the course of 120 days. Solid lines represent AmbT plants while dashed lines represent ElevT. No ElevT mature tubers were collected at 90d, so no data is shown for that point. TPM values were determined from *Salmon* (Patro et al. 2017) and averaged per biological replicate (*n* = 1 or 2) (*p* < 0.10 = *; *p* < 0.05 = **).

**Supplementary Figure 5E.**
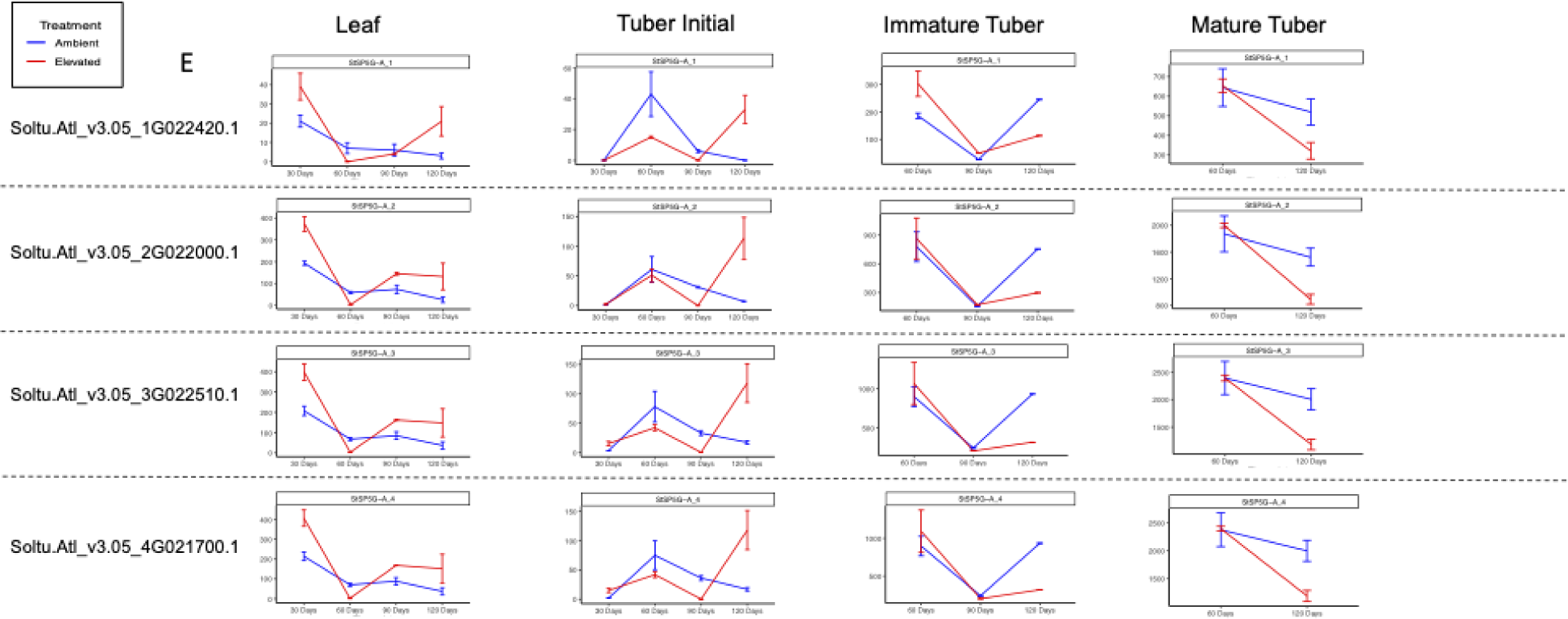
TPMs of known tuberization inhibitor genes *StSP5G-A* syntelogs in each tissue type over the course of 120 days. Solid lines represent AmbT plants while dashed lines represent ElevT. No ElevT mature tubers were collected at 90d, so no data is shown for that point. TPM values were determined from *Salmon* (Patro et al. 2017) and averaged per biological replicate (*n* = 1 or 2) (*p* < 0.10 = *; *p* < 0.05 = **).

**Supplementary Figure 5F.**
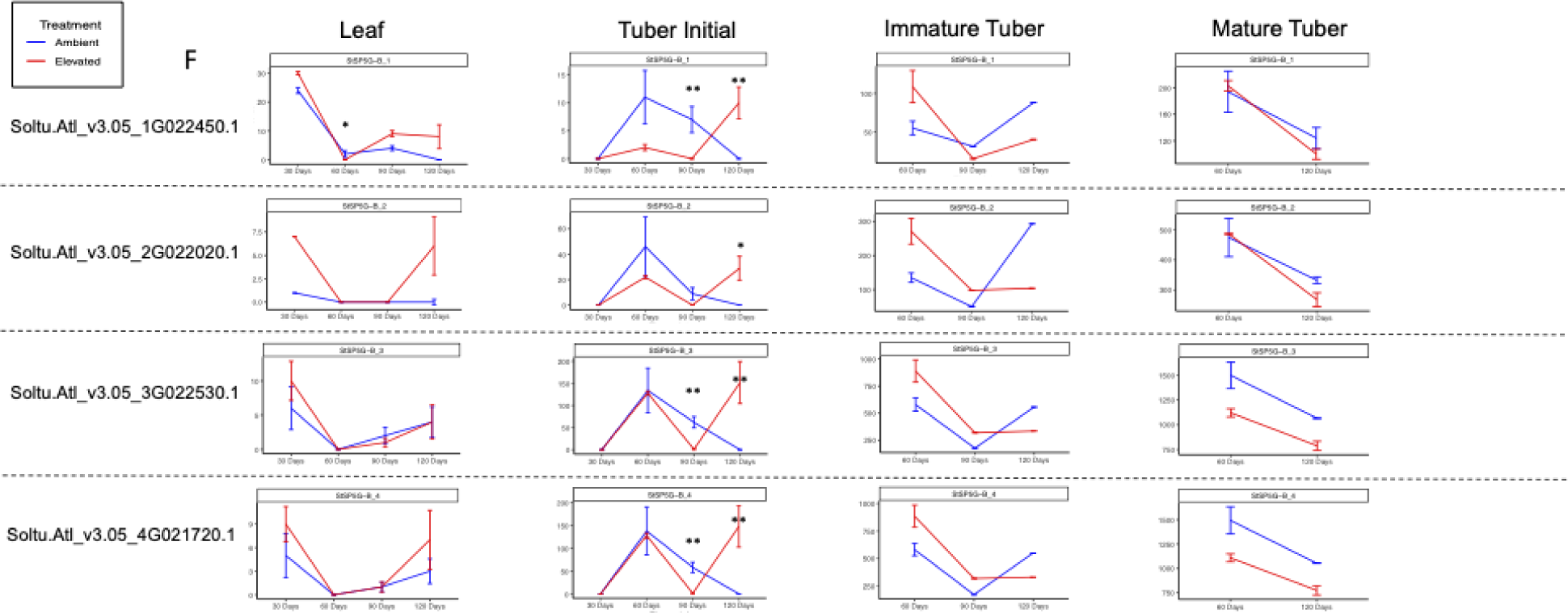
TPMs of known tuberization inhibitor genes *StSP5G-B* syntelogs in each tissue type over the course of 120 days. Solid lines represent AmbT plants while dashed lines represent ElevT. No ElevT mature tubers were collected at 90d, so no data is shown for that point. TPM values were determined from *Salmon* (Patro et al. 2017) and averaged per biological replicate (*n* = 1 or 2) (*p* < 0.10 = *; *p* < 0.05 = **).

**Supplementary Figure 5G.**
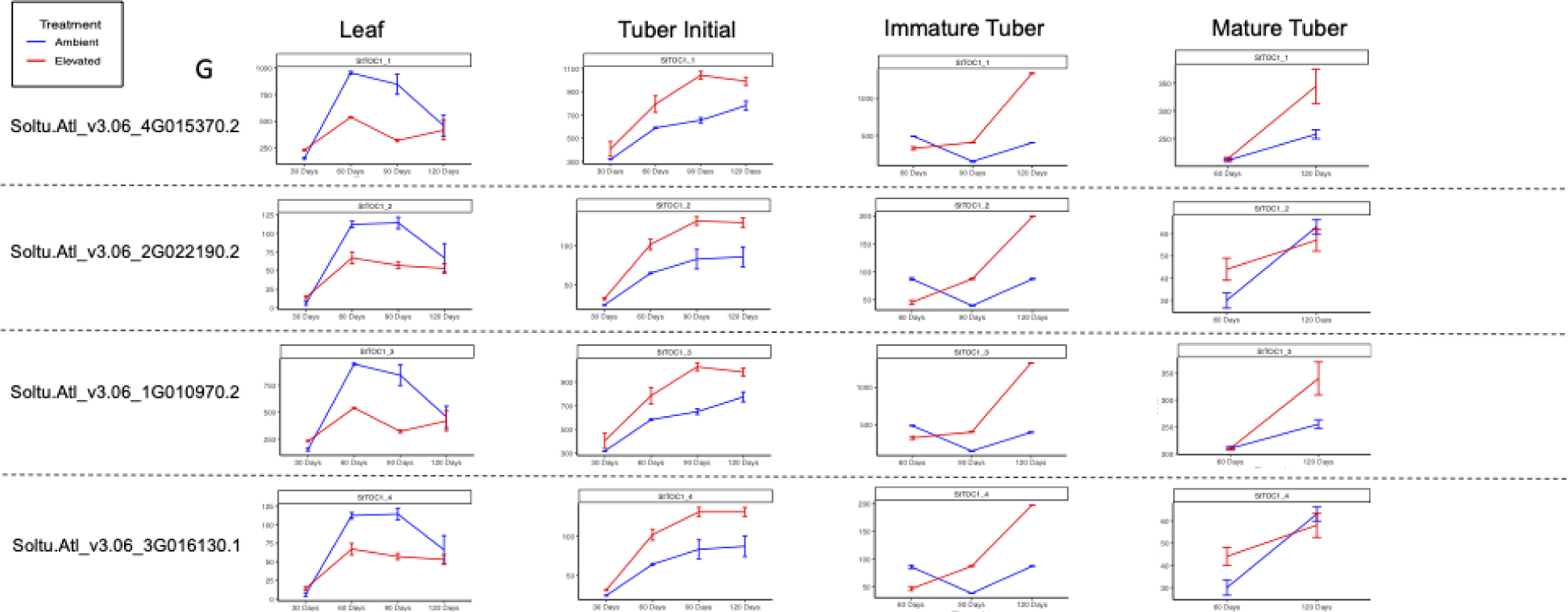
TPMs of known tuberization inhibitor gene *StTOC1* syntelogs in each tissue type over the course of 120 days. Solid lines represent AmbT plants while dashed lines represent ElevT. No ElevT mature tubers were collected at 90d, so no data is shown for that point. TPM values were determined from *Salmon* (Patro et al. 2017) and averaged per biological replicate (*n* = 1 or 2) (*p* < 0.10 = *; *p* < 0.05 = **).

